# CRISPR-GPT for Agentic Automation of Gene Editing Experiments

**DOI:** 10.1101/2024.04.25.591003

**Authors:** Yuanhao Qu, Kaixuan Huang, Ming Yin, Kanghong Zhan, Dyllan Liu, Di Yin, Henry C. Cousins, William A. Johnson, Xiaotong Wang, Mihir Shah, Russ B. Altman, Denny Zhou, Mengdi Wang, Le Cong

**Author notes:** Corresponding authors, (M.W.), (L.C.). These authors contributed equally (Yuanhao Qu, Kaixuan Huang).

## Abstract

Performing effective gene-editing experiments requires a deep understanding of both the CRISPR technology and the biological system involved. Meanwhile, despite their versatility and promise, Large Language Models (LLMs) often lack domain-specific knowledge and struggle to accurately solve biological design problems. We present CRISPR-GPT, an LLM agent system to automate and enhance CRISPR-based gene-editing design and data analysis. CRISPR-GPT leverages the reasoning capabilities of LLMs for complex task decomposition, decision-making, and interactive human-artificial intelligence (AI) collaboration. This system incorporates domain expertise, retrieval techniques, external tools, and a specialized LLM fine-tuned with open-forum discussions among scientists. CRISPR-GPT assists users in selecting CRISPR systems, experiment planning, designing gRNAs, choosing delivery methods, drafting protocols, designing assays, and analyzing data. We showcase the potential of CRISPR-GPT by knocking-out four genes with CRISPR-Cas12a in a human lung adenocarcinoma cell line and epigenetically activating two genes using CRISPR-dCas9 in human melanoma cell line. CRISPR-GPT enables fully AI-guided gene-editing experiment design and analysis across different modalities, validating its effectiveness as an AI co-pilot in genome engineering.

## Main text

Large language models (LLMs) have demonstrated exceptional capabilities in language skills and encapsulate a substantial amount of world knowledge^19–23^. Recent research has also enhanced LLMs with external tools, improving their problem-solving abilities and efficiencies^24–26^. Moreover, LLMs have also demonstrated potential as tool makers^27^ and black-box optimizers^28^. To this end, researchers have explored LLM-based specialized models for various scientific domains^29,30^, particularly for mathematics and chemistry tasks. ChemCrow^31^ uses tool-augmented LLM for solving a range of chemistry-related tasks such as paracetamol synthesis, whereas Co-scientist^32^ integrated automated experimentation, achieving successful optimization of palladium-catalyzed cross-coupling reaction. LLMs have also shown initial promise in generating biological protocols, as demonstrated by studies like BioPlanner^78^. While recent advancements, such as OpenAI’s o1 preview, have improved reasoning abilities in areas like mathematics and coding, progress in biological tasks remains comparatively limited. This limitation stems from general-purpose LLMs’ lack of in-depth understanding of biology, compounded by the unique challenges of biological experiments, including the variability of living systems, the noisy nature of biological data, and the highly specialized, less transferable nature of biological skills and tools.

Gene editing has transformed biological research and medicine, allowing for precise DNA modifications for both therapeutic and experimental applications. CRISPR-Cas, the most well-known gene-editing technology, originated from bacterial immune systems^1–9^. Its development has led to advanced techniques like CRISPR activation and interference (CRISPRa/i)^12–16^, base-editing^17,18^, and prime-editing^10,11^, creating a powerful toolkit for genetic modification and epigenetic modulation. In basic biomedical research, CRISPR gene-editing has become one of the most frequently used laboratory techniques: at the largest non-profit plasmid DNA repository, Addgene, 8 of the 15 top requested plasmids worldwide were for CRISPR gene-editing^73^. On the application side, CRISPR has produced the first permanent cure for Sickle Cell Disease (SCD)^74^ and β-thalassemia^75^, as well as facilitating plant engineering for sustainable agriculture^5^. As one of the most powerful biotechnologies, numerous software and protocols exist for specific gene-editing tasks. Despite these resources, an end-to-end solution—from CRISPR-Cas system selection, gRNA design, off-target evaluation, to delivery and data analysis—remains complex, particularly for newcomers. AI-assisted tools can simplify gene-editing experiment design and data analysis, making the technology more accessible and accelerating scientific and therapeutic discoveries.

We introduce CRISPR-GPT, a solution that combines the strengths of LLMs with domain-specific knowledge, chain of thought reasoning, instruction finetuning, retrieval techniques and tools. CRISPR-GPT is centered around LLM-powered planning and execution agents (**Figure 1**). This system leverages the reasoning abilities of general-purpose LLMs and multi-agent collaboration for task decomposition, constructing state machines, and automated decision-making (**Figure 2a**). It draws upon expert knowledge from leading practitioners and peer-reviewed published literatures in gene editing for retrieval-augmented generation (RAG)^31^.

**Fig. 1.**
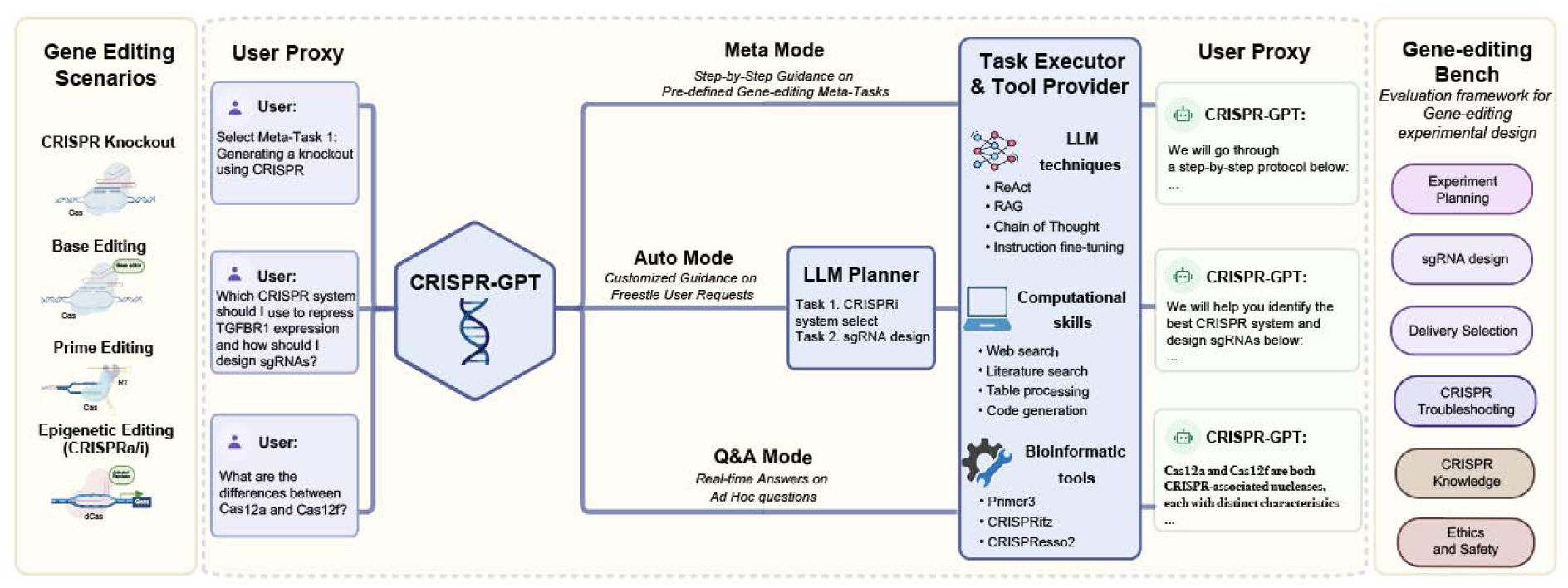
Overview of CRISPR-GPT. CRISPR-GPT is an LLM-powered multi-agent system designed to provide AI copiloting for human researchers in gene-editing. It supports four primary gene-editing modalities: knockout, base editing, prime editing, and epigenetic editing (CRISPRa/i). The system offers three user interaction modes—Meta Mode (step-by-step guidance on predefined tasks), Auto Mode (customized guidance based on user requests), and QA Mode (real-time answers to ad hoc questions)—to streamline experiment design and planning. CRISPR-GPT consists of four core components: the User Proxy, LLM Planner, Task Executor, and Tool Provider. Together, these components are equipped with a comprehensive suite of tools and decision-support capabilities to facilitate the design, planning, and analysis of gene-editing workflows. To evaluate CRISPR-GPT’s performance, we developed the Gene-editing Bench, a framework of 288 test cases covering tasks such as experimental planning, sgRNA design, delivery method selection, and more. Panel was originally created in BioRender. Qu, Y. (2025) https://BioRender.com/tb8sq6f.

**Fig. 2.**
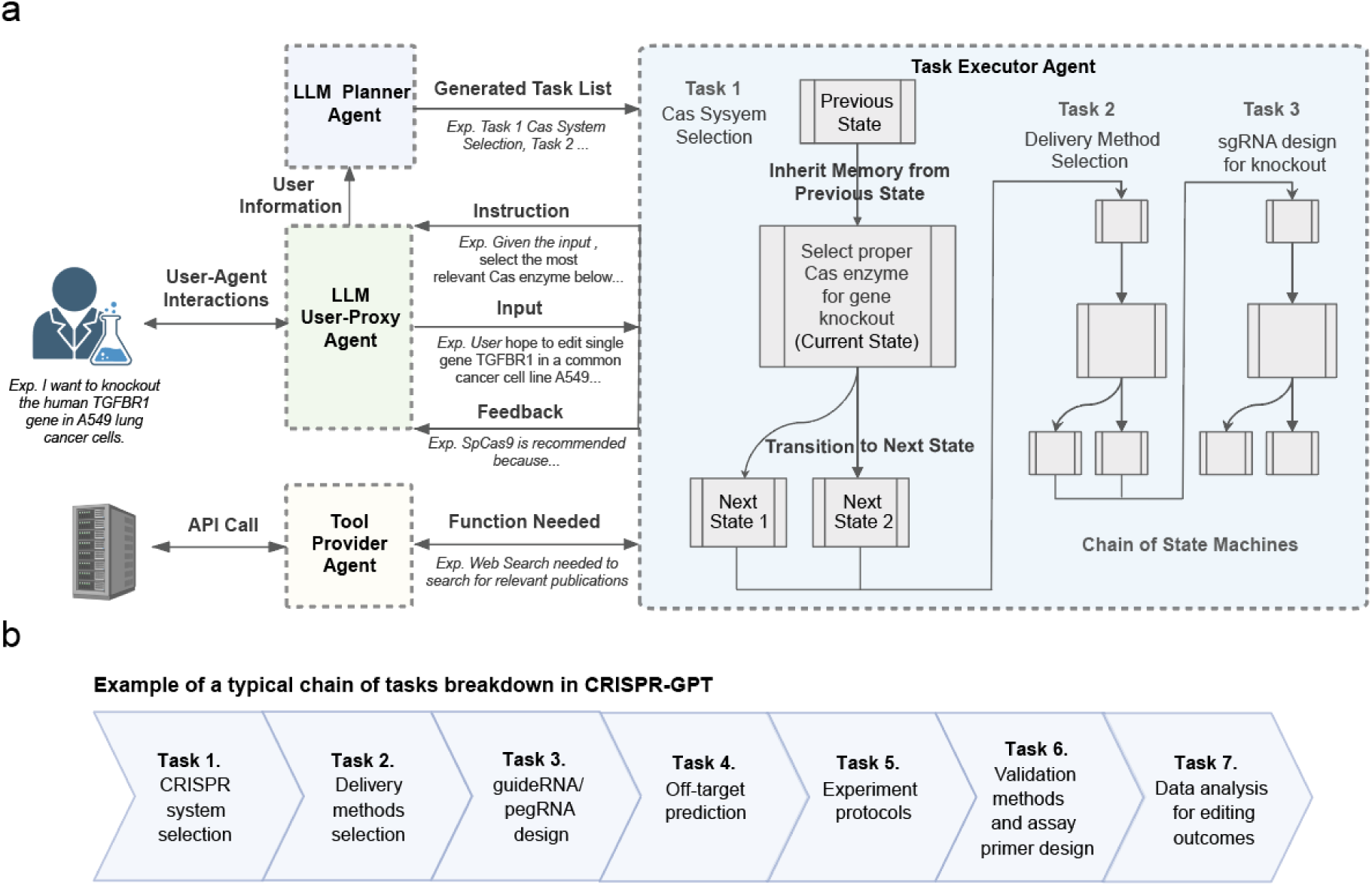
CRISPR-GPT adopts a compositional, multi-agent architecture to enable human-AI collaboration and automated experimental designs. **a,** The backbone of CRISPR-GPT involves multi-agent collaboration between four core components: (1) *LLM Planner Agent* is responsible for configuring tasks based on the user’s needs. It automatically performs task decomposition based on the user’s request, the descriptions of the currently supported tasks, and internal knowledge. The state machines of the selected tasks are chained together to fulfill the user’s request. (2) *Task Executor Agent* implements the chain of state machines from the Planner Agent, and is responsible for providing instructions and feedback, receiving input from User-Proxy Agent, and calling external tools. State machines are central to the Task Executor, where each state is responsible for one round of interaction with the user. The instruction is provided to the user first with sufficient information for the current decision-making step and the required inputs. After receiving the response from the user, it provides output and feedback, where Tool Providers are potentially called during the execution of the state. Afterward, the state machine transits to the next state. (3) *LLM User-Proxy Agent* is responsible for interacting with the Task Executor on behalf of the user, where the user can monitor the process and provide corrections to the User-Proxy Agent if the generated content needs modification or improvement. It generates responses to every step of the state machine on behalf of the user. (4) Tool Providers support diverse external tools and connect to search engines or databases via API calls Part of the panel was created in BioRender. Qu, Y. (2025) https://BioRender.com/svkmgjk. **b.** Breakdown of individual tasks in a typical CRISPR-GPT workflow for gene-editing experiments.

## Results

### Building AI co-pilot harnessing LLM’s reasoning ability

CRISPR-GPT supports four major gene-editing modalities and 22 gene-editing experiment tasks (**Figure 1, Supp. Table 1**). It offers tunable levels of automation via three modes: Meta, Auto, and QA. They are designed to accommodate users from novice PhD-level scientist fresh to gene-editing, to domain experts looking for more efficient, automated solutions for selected tasks (**Figure 1**). “Meta Mode” is designed for beginner researchers, guiding them through a sequence of essential tasks from selection of CRISPR systems, delivery methods, to designing gRNA, assessing off-target efficiency, generating experiment protocols and data analysis. Throughout this decision-making process, CRISPR-GPT interacts with users at every step, provides instructions, and seeks clarifications when needed. “Auto Mode’’ caters to advanced researchers and does not adhere to a predefined task order. Users submit a free-style request, and the LLM-planner decomposes this into tasks, manages their interdependence, builds a customized workflow and executes them automatically. It fills in missing information based on initial inputs and explains its decisions and thought process, allowing users to monitor and adjust the process. “Q&A Mode” supports users with on-demand scientific inquiries about gene-editing.

To assess the AI agent’s capabilities to perform gene-editing research, we compiled an evaluation testset, Gene-editing-Bench, from both public sources and human experts (details in **Supp. Note C**). This testset covers a variety of gene-editing tasks (**Figure 1**). By using the testset, we performed extensive evaluation of CRISPR-GPT’s capabilities in major gene-editing research tasks, such as experiment planning, delivery selection, sgRNA design, and experiment troubleshooting. Additionally, we invited human experts to perform a thorough user experience evaluation of CRISPR-GPT and collected valuable human feedback.

Further, we implement CRISPR-GPT in real-world wet labs. Using CRISPR-GPT as an AI co-pilot, we demonstrate a fully AI-guided knockout of four genes—*TGFBR1*, *SNAI1*, *BAX*, and *BCL2L1*—using CRISPR-Cas12a in human lung adenocarcinoma cell line, as well as AI-guided CRISPR-dCas9 epigenetic activation of two genes—*NCR3LG1, CEACAM1*—in a human melanoma model. All these wet-lab experiments were carried by junior researchers not familiar with gene-editing. They both succeeded on the first attempt, confirmed by not only editing efficiencies, but also biologically relevant phenotypes and protein-level validation, highlighting the potential of LLM-guided biological research.

CRISPR-GPT is a multi-agent, compositional system involving a team of LLM-based agents, including an LLM Planner Agent, a User-Proxy Agent, Task Executor Agents, and Tool Provider Agents (**Figure 2a**). These components are powered by LLMs to interact with one another as well as the human user. We also refer to the full system as an “agent” to encapsulate the overall functionalities.

To automate biological experiment design and analysis, we view the overall problem as sequential decision-making. This perspective frames the interaction between the user and the automated system as a series of decision-making steps, each essential for progressing towards the ultimate goal. Take the Auto Mode for example. A user can initiate the process with a meta-request, for example, “I want to knock out the human TGFBR1 gene in A549 lung cancer cells”. In response, the agent’s LLM planner will analyze the user’s request, drawing on its extensive internal knowledge base via retrieval techniques. Leveraging the reasoning abilities of the base LLM, the planner generates a chain-of-thought^44^ reasoning path and chooses an optimal action from a set of plausible ones, while following expert-written guidelines. Consequently, the Planner breaks down the user’s request into a sequence of discrete tasks, for example “CRISPR/Cas system selection” and “gRNA design for knockout”, while managing inter-dependencies among these tasks. Each individual task is solved by an LLM-powered state machine, via the Task Executor, entailing a sequence of states to progress towards the specific goal. After the meta-task decomposition, the Task Executor will chain the state machines of the corresponding tasks together into a larger state machine and begin the execution process, systematically addressing each task in sequence to ensure the experiment’s objectives are met efficiently and effectively (**Figure 2a**).

The User-Proxy Agent is responsible for guiding the user throughout the decision-making process via multiple rounds of textual interactions (typical user interactions required by each task detailed in Supp. Table 2). At each decision point, the internal state machine presents a “state variable” to the User-proxy Agent, which includes the current task instructions and specifies any necessary input from the user to proceed. The user-proxy agent then interprets this state given the user interactions and makes informed decisions as input to Task Executor on behalf of the user. Subsequently, the User-Proxy Agent receives feedback from the Task Executor, including the task results and the reasoning process that led to those outcomes. Concurrently, the user-proxy agent continues to interact with the user and provides her with instructions, continuously integrating her feedback to ensure alignment with the user’s objectives (detailed in Method and Figure 2a, Supp. Fig.1,).

To enhance the LLM with domain knowledge, we enable the CRISPR agent to retrieve and synthesize information from published protocols, peer-reviewed research papers, expert-written guidelines, and to utilize external tools and conduct web searches via Tool Provider Agents (**Figure 2a**).

For an end-to-end gene-editing workflow, CRISPR-GPT typically constructs a chain of tasks that includes selecting the appropriate CRISPR system, recommending delivery methods, designing gRNAs, predicting off-target effects, selecting experimental protocols, planning validation assays, and performing data analysis **(Figure 2b)**. The system’s modular architecture facilitates easy integration of additional functionalities and tools. CRISPR-GPT serves as a prototype LLM-powered AI co-pilot for scientific research, with potential applications extending beyond gene editing.

### CRISPR-GPT agents automate gene-editing research tasks

CRISPR-GPT is able to automate gene-editing research via several key functionalities. For each functionality we discuss the agentic implementation and evaluation results.

#### Experiment Planning

The Task Planner Agent is charged with directing the entire workflow and breaking down the user’s meta-request into a task chain (**Figure 2b**). While the Planner selects and follows a predefined workflow in the Meta Mode, it is able to intake free-style user requests and auto-build a customized workflow in the Auto Mode. For example, a user may only need part of the predesigned workflow including CRISPR/Cas system selection, delivery method selection, guideRNA design and experimental protocol selection before the experiment. Then the Task Planner Agent extracts the right information from the user request and assembles a customized workflow to suit user needs (**Figure 3a**). To evaluate CRISPR-GPT’s ability to correctly layout gene-editing tasks and manage inter-task dependence, we compiled a planning testset, as a part of the Gene-editing-Bench, with user requests and golden answers curated by human experts. Using this testset, we evaluated CRISPR-GPT in comparison with prompted general LLMs, showing that CRISPR-GPT outperforms general LLMs in planning gene-editing tasks (**Figure 3b**). The CRISPR-GPT agent driven by GPT-4o scored over 0.99 in accuracy, precision, recall, F1 score, and had less than 0.05 in the normalized Levenshtein distance between agent-generated plans and golden answers (**Figure 3b**). For extensive description of the testset and evaluation, please see **Supp. Note C1.**

**Fig. 3.**
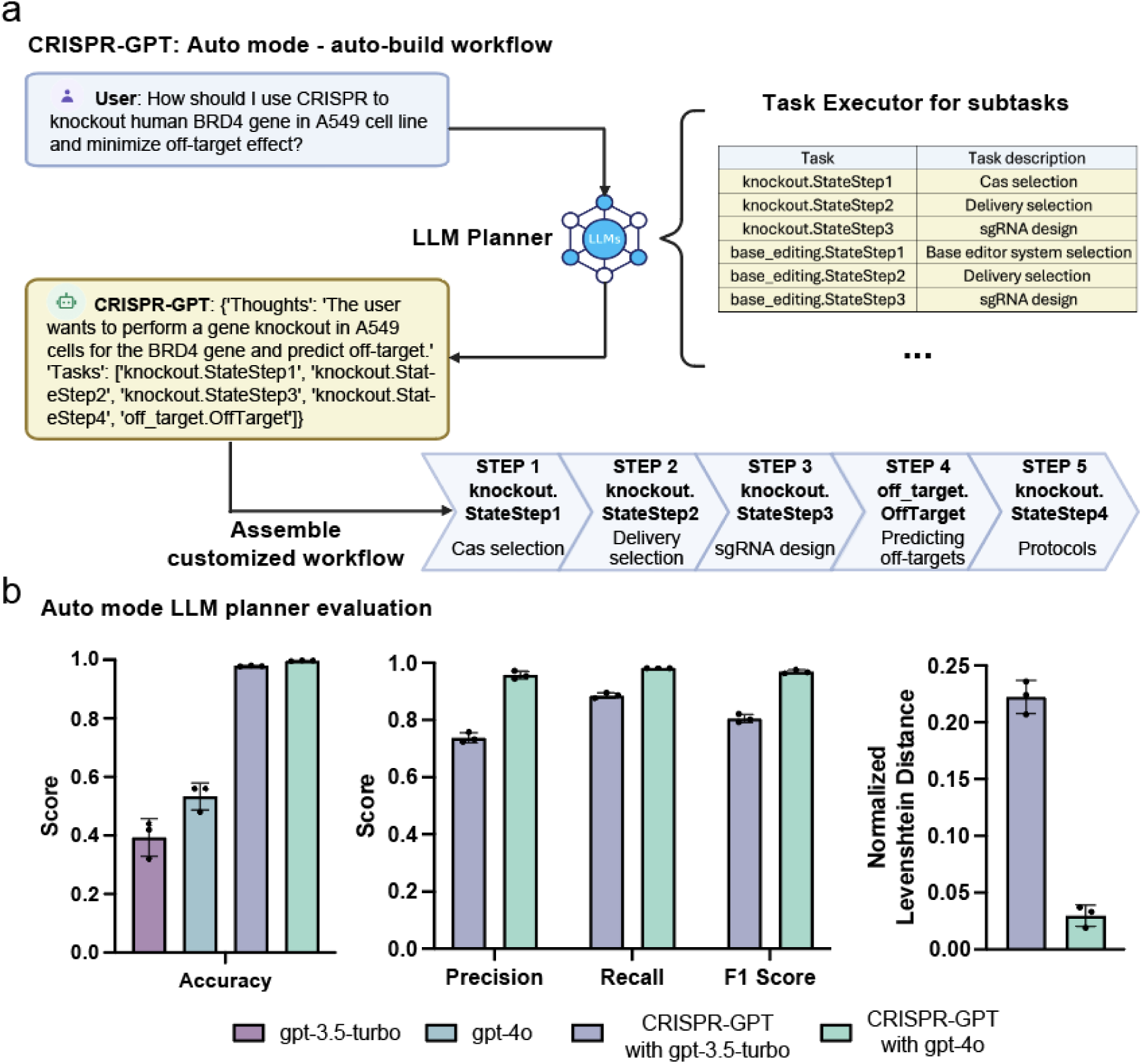
Task decomposition and experiment planning in CRISPR-GPT Auto-mode with performance evaluation. **a,** The LLM Planner Agent automatically breaks down the user’s meta-request to a sequence of tasks. Then it assembles a customized workflow of the chained tasks to meet the user’s needs. Part of the panel was created in BioRender. Qu, Y. (2025) https://BioRender.com/qy4v. **b,** Evaluation of the LLM Planner using a gene-editing planning testset. For each test case, we generate three independent answers from each model and report the average scores (see **Supp. Note C**). Data shown are the mean ± s.d. (*n* = 3 per group).

#### Delivery Method Selection

We present and evaluate the delivery agent of CRISPR-GPT (**Figure 4a-b**). Delivery is a critical step for all gene-editing experiments. CRISPR-GPT equips LLM with expert-tailored instructions and external tools to choose delivery methods. Specifically, the agent first tries to understand the biological system that the user is planning to edit. It extracts keywords for the target cell/tissues/organisms, performs Google search, and summarizes the results. Then, given its own knowledge and search results, CRISPR-GPT matches the user case with a major biological category–cell lines, primary cells, in vivo, etc.– which reduces the possible options to a focused set of candidate methods. Next, CRISPR-GPT performs literature search with user and method-specific keywords, and ranks the candidate methods based on citation metrics to suggest a primary and a secondary delivery method (**Figure 4a**). To evaluate the performance of this module, we compiled test cases including 50 biological systems as a part of the Gene-editing-Bench. For each case, we invited three human experts to score potential delivery options and utilized those as ground-truth. We then evaluated the output of CRISPR-GPT and baseline models by comparing to the pre-compiled ground-truth score sheet. We found that CRISPR-GPT outperforms the baseline gpt-4, gpt-3-turbo models (**Figure 4b**). The agent has a substantial edge on difficult tasks such as those involving hard-to-transfect cell lines and primary cell types. We also noticed that including an additional literature search step improves the agent’s performance only moderately. More details about the delivery selection evaluation can be found in **Supp. Note C2.**

**Fig. 4.**
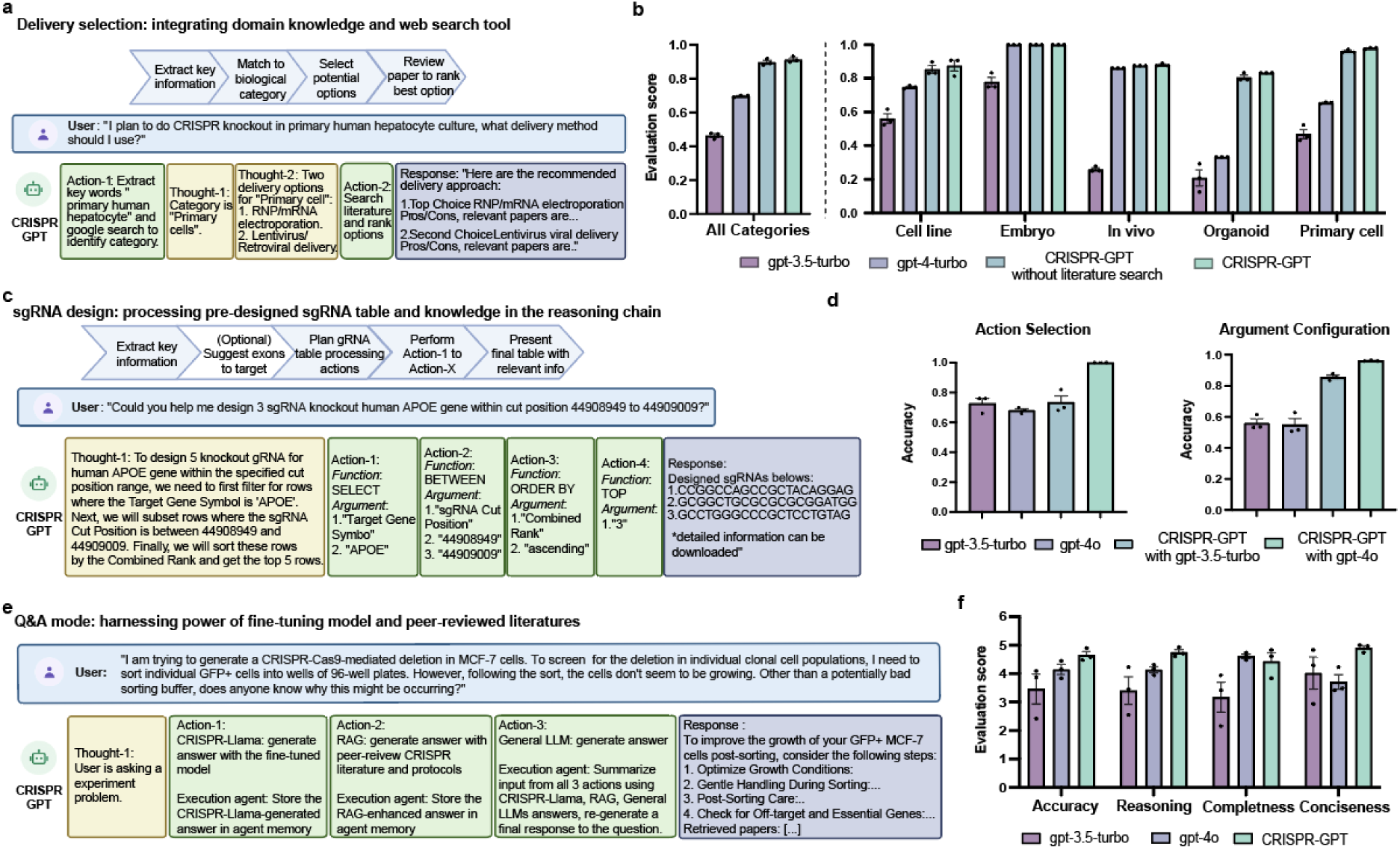
CRISPR-GPT automates gene-editing research and experiment tasks. **a,** Design of delivery method selection agent in CRISPR-GPT, showing the workflow, example request, and a series of agent thoughts-actions to identify most suitable delivery methods for the user’s needs. **b,** Evaluation results of delivery method selection using CRISPR-GPT and baseline models Data shown are the mean ± s.d. (*n* = 3 per group).. **c,** Design of guideRNA design agent in CRISPR-GPT, showing the workflow, example request, and a series of agent thoughts-actions to select top-ranked gRNA customized to user’s request. **d,** Evaluation results of gRNA design using CRISPR-GPT and baseline models. Models were prompted to generate functions and associated parameters to design gRNAs requested by the user. Data shown are the mean ± s.d. (*n* = 3 per group). **e,** Design of QA Mode in CRISPR-GPT, showing the workflow, example request, and a series of agent thoughts-actions to answer gene-editing questions. **f,** Evaluation of CRISPR-GPT and baseline models for answering gene-editing research questions. Models were prompted to generate answers, which were anonymized, evaluated by three human experts in a fully blind set-up. Scores range from 1 (lowest) to 5 (highest). All scores from above were from three independent trials (details on evaluation in **Supp. Note C**). Data shown are the mean ± s.d. (*n* = 3 per group).

#### guideRNA design

Good guide RNA (gRNA) design is crucial for the success of CRISPR experiments. Various gRNA design tools and softwares, such as CRISPick^33–36^ and ChopChop^79^, are available. However, we believe there are two key challenges in general usage: 1. Choosing a trustworthy source. 2. Difficulty in quickly identifying gRNAs that suit specific user requirements or experiment contexts, often requiring lengthy sorting, ranking, or literature review. To address these issues, we utilized pre-designed gRNA tables from CRISPick, a reputable and widely used tool. We leverage the reasoning capabilities of LLMs to accurately identify regions of interest, and quickly extract relevant gRNAs. This approach is similar to the recently proposed “chain-of-tables” methodology^77^ (**Figure 4c, Ext. Data Fig. 1a, Supp. Demo Video 1,2**). To evaluate the ability of CRISPR-GPT to correctly retrieve gRNAs, we compiled a gRNA design test set with ground truth from human experts (detailed in **Supp. Note C3**). CRISPR-GPT agent outperforms the baseline LLMs, in accurately selecting gRNA design actions and configuring the arguments (**Figure 4d**).

Further, we picked a real-world test case from a cancer biology study, in which many highly-ranked gRNA designs did not generate biological phenotypes, even when their editing efficiencies were high^76^. Instead, the authors of the study had to design gRNAs manually against Exons encoding important functional domains within a gene, and Exon-selected gRNAs induced expected cancer-killing effects. We tested CRISPR-GPT for designing gRNAs targeting BRD4 gene from this study, and compared results with those generated by CRISPick and CHOPCHOP (**Ext. Data Figure 1**). CRISPR-GPT was uniquely able to select the key exons, Exon3-4, within BRD4. In contrast, gRNAs designed by CRISPick or CHOPCHOP would be likely ineffective, as 7 out of 8 gRNAs mapped to non-essential Exons (**Ext. Data Figure 1**). Taken together, our results support the benefit and validity of this module.

#### Other Functions and Tools

CRISPR-GPT provides specific suggestions on the choice of the CRISPR system, experimental and validation protocol selection, by leveraging LLM’s reasoning ability and retrieving information from an expert-reviewed knowledge base. It also offers automated gRNA off-target prediction, primer design for validation experiments, and data analysis. In particular, the agent provides fully automated solutions to run external softwares, such as Primer3^42^, CRISPRitz^50^ and CRISPResso2^62^ (**Supp. Table 1**). We focused on implementing these tools as they are considered gold-standard in respective tasks, and have been extensively validated in prior work.

### Problem-solving via fine-tuning LLM on scientific discussion

General-purpose LLMs may possess broad knowledge but often lack the deep understanding of science needed to solve research problems. To enhance the CRISPR-GPT agent’s capacity in answering advanced research questions, we build a QA Mode that synthesizes information from multiple resources, including published literature, established protocols, and discussions between human scientists, utilizing a combination of RAG technique, a fine-tuned specialized model and a general LLM (for which we picked gpt-4o). (**Methods**).

To enhance the QA mode’s capacity to “think” like a scientist for problem solving, we sought to train a specialized language model using real scientific discussions among domain experts. The fine-tuned model is used as one of the multiple sources of knowledge for the QA mode (**Figure 4e)**. To this end, we collected 11 years of open-forum discussions from a public Google Discussion Group on CRISPR gene-editing, starting from 2013 (**Supp. Note B**). The discussion group involved a diverse cohort of scientists worldwide. This dataset, comprising approximately 4,000 discussion threads, was curated into an instructional dataset with over 3,000 question-and-answer pairs (**Supp. Note B**). Using this dataset, we fine-tuned an 8-billion-parameter LLM based on the Llama3-instruct model^59^. The fine-tuned model, which we call CRISPR-Llama3, has improved abilities in gene-editing questions, outperforms the baseline model on basic questions by a moderate 8% and on real-world research questions by ∼20% (**Supp.** Fig. 2-3). We integrate this fine-tuned LLM into the QA Mode as a “brainstorming source”, enabling the agent to generate ideas like a human scientist and provide a second opinion for difficult queries (**Figure 4e**).

To assess the performance of the QA Mode, we used the Gene-editing-Bench QA testset (**Supp. Note C**). The test questions encompass basic gene-editing knowledge, experimental troubleshooting, CRISPR application in various biological systems, ethics and safety. We prompted CRISPR-GPT, gpt-3.5-turbo, and gpt-4o to generate responses to test questions. Three human experts scored the answers in a fully blinded setting. The test demonstrated that the QA Mode outperformed baseline LLMs in accuracy, reasoning, and conciseness, with improvement of 12%, 15%, and 32%, respectively, versus GPT-4o (**Figure 4f**). Human evaluators observe that general-purpose LLMs sometimes make factual errors and tend to provide extensive answers not all relevant to the questions. For example, one question is about solving cell growth issues in an experiment where a scientist performed Cas9 editing followed by single-cell sorting using MCF-7 cells. For this question, the QA Mode provided a concise, accurate summary of potential reasons and actionable solutions. In contrast, GPT-4o responded with a long list of 9 factors/options, but at least 2 of them are not applicable to MCF-7 cells (**Ext. Data Figure 2**). This, and other examples (**Ext. Data Figures. 3-4**) showcase the advantage of CRISPR-GPT QA Mode. Overall, evaluation results confirmed that the multi-source QA Mode is better at answering advanced research questions about gene-editing.

#### CRISPR-GPT excels in human-AI collaboration

To further evaluate the human user experience of CRISPR-GPT, we assembled a panel of eight gene-editing experts to assess the agents’ performance for end-to-end experiment covering all 22 individual tasks (See Supp. Table 2 and Supp. Table 3 demos). The experts were asked to rate their experiences in four dimensions: Accuracy, Reasoning and Action, Completeness, and Conciseness (see **Supp. Note C** for detailed rubrics). CRISPR-GPT demonstrated improved accuracy and strong capabilities in reasoning and action, whereas general LLMs, such as GPT-4o, often included errors and were prone to hallucination (**Figure 5a, b)**.

**Fig. 5.**
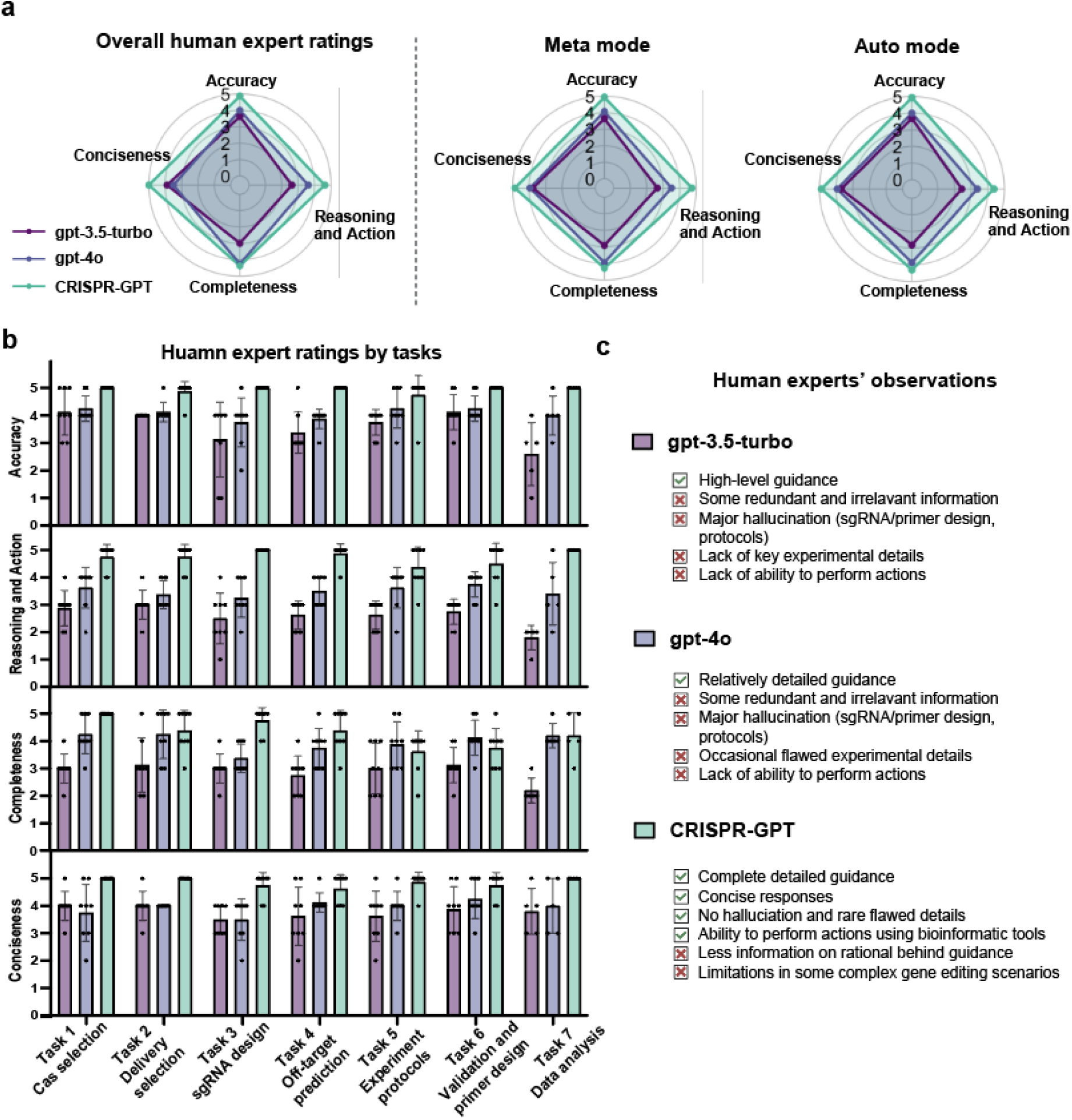
CRISPR-GPT outperforms general-purpose LLM for gene-editing research in human user experiences. **a,** Human user experience: evaluation of CRISPR-GPT for end-to-end gene-editing copiloting. Human experts scored performances from 1 (lowest) to 5 (highest). See detailed procedure and rubrics in **Supp. Note C** (Full chat history and video demo listed in **Supp. Table 2**). **b,** Human user experience: evaluation results breakdown by major gene-editing tasks. Data shown are the mean ± s.d. (*n* = 8 per group). **c,** User observations on the strengths and limitations of CRISPR-GPT compared to baseline LLMs.

Highlighted by human evaluators’ observations (**Figure 5c**), the CRISPR-GPT agent provides users with more accurate, concise, and well-rounded instructions to execute the planned experiments. The ability of CRISPR-GPT to perform specialty gene-editing tasks, such as exon-selected gRNA design, customized off-target prediction, and automated sequencing data analysis, reinforced its advantage versus general-purpose LLMs. This is confirmed by the task-specific evaluation results (**Figure 5b**). Despite its strengths, CRISPR-GPT struggled with complex requests and rare biological cases, highlighting areas for improvement (limitations in **Supp. Note D**).

#### Real-world demonstration for fully AI-guided gene-editing

To showcase and validate CRISPR-GPT’s ability as an AI co-pilot to general biologists, we enlisted two junior researchers unfamiliar with gene-editing. They used CRISPR-GPT in two real-world experiments: to design and conduct a multi-gene knockout and an epigenetic editing experiment, respectively, from scratch.

In the first experiment, the junior researcher conducted gene knockouts in the human A549 lung adenocarcinoma cell line, targeting four genes involved in tumor survival and metastasis: TGFBR1, SNAI1, BAX, and BCL2L1 (**Figure 6**). The experiment was designed from scratch with CRISPR-GPT (**Figure 6a**). Based on user-AI interaction, enAsCas12a was selected for its multi-target editing capability and low off-target effects. For delivery, CRISPR-GPT recommended lentiviral transduction for stable Cas and gRNA expression. The gRNAs for the four target genes were designed through CRISPR-GPT. Furthermore, CRISPR-GPT provided step-by-step protocols for gRNA cloning, lentivirus production, and viral delivery into A549 cells. To validate the editing, the researcher followed CRISPR-GPT’s NGS protocol, using assay primers designed via the integrated Primer3 tool. After generating the NGS data, the raw sequencing files were uploaded into CRISPR-GPT for automated analysis through the CRISPResso2 pipeline. The analysis reports, sent directly via email, summarized the editing outcomes and showed consistently ∼80% high efficiency across all target genes (**Figure 6b**, **Supp. Demo Video 3**, user interactions summarized in **Supp. Table 4**, full chat history listed in **Supp. Table 2**). To further assess the biological phenotypes of TGFBR1, SNAI1 knockout in A549 cells, the researcher conducted an Epithelial-mesenchymal transition (EMT) induction experiment by treating A549 cells with TGFβ (**Figure 6c**, and Methods). The qPCR results revealed that the knockout A549 cell lines (A549 TGFBR1 KO and A549 SNAI1 KO) showed up to 9-fold reduction in CDH1 expression change, and up to 34-fold reduction in VIM expression change, which are both key marker genes in the EMT process. This confirms the biological role of TGFBR1 and SNAI1 signaling in driving EMT progression (a crucial driver of metastasis) in lung cancer cells (**Figure 6d**).

**Fig. 6.**
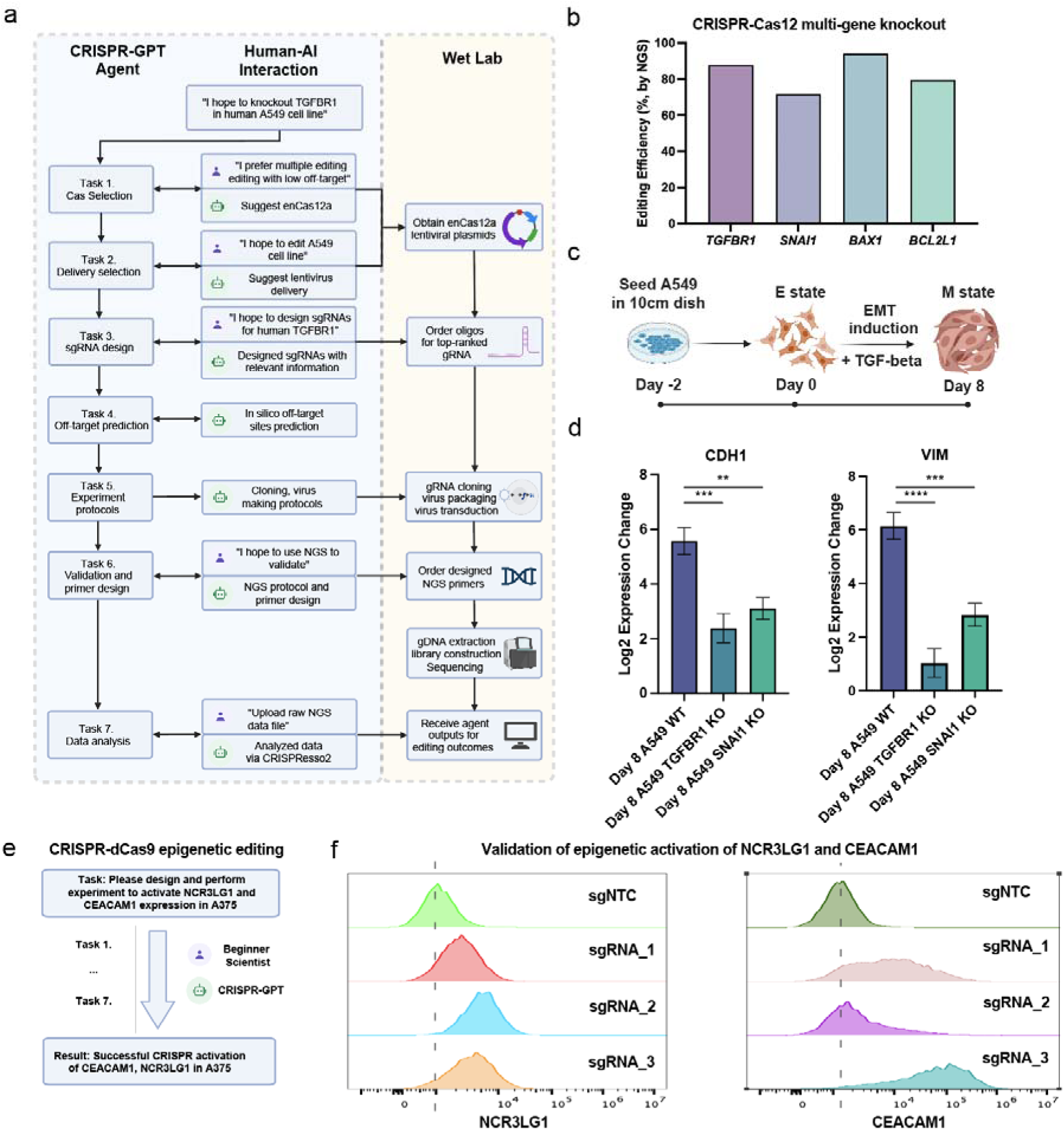
Wet-lab demonstrations of CRISPR-GPT in knockout and activation experiments. **a,** The full workflow of CRISPR-GPT-guided knockout experiment of *TGFBR1*, *SNAI1*, *BAX1*, and *BCL2L1* through multiple rounds of human-AI interaction (*TGFBR1* knockout is shown as an example, see **Supp. Demo Video 3** and full chat history listed in **Supp. Table 2**). Panel was originally created in BioRender. Qu, Y. (2025) https://BioRender.com/iim1m2t and processed. **b,** Editing efficiencies for *TGFBR1*, *SNAI1*, *BAX1*, and *BCL2L1* measured via next-generation sequencing, analyzed using CRISPResso2 and CRISPR-GPT. **c,** Schematic of the EMT induction process via TGF-β treatment (see Methods). Panel was originally created in BioRender. Qu, Y. (2025) https://BioRender.com/iim1m2t and processed. **d,** Functional outcomes of *TGFBR1* and *SNAI1* knockout in A549 cells after EMT induction by TGF-β. qPCR analysis shows reduced expression changes in EMT marker genes (*CDH1*, *VIM*) in A549 *TGFBR1* and *SNAI1* knockout cells compared with A549 Wildtype (WT) cells, confirming successful knockouts of *TGFBR1*/*SNAI1*. Data shown are the mean ± s.d. (*n* = 3 per group). One-way ANOVA was used to calculate the statistics. ****, p<0.0001. **e,** Simplified workflow of a beginner researcher activating *NCR3LG1* and *CEACAM1* expression through multi-round interactions with CRISPR-GPT (full chat history listed in **Supp. Table 2**). **f,** Editing outcomes of *NCR3LG1* and *CEACAM1* activation using CRISPR-GPT designed sgRNAs, measured via flow cytometry (see Methods).

In the second experiment, the junior researcher performed epigenetic editing to activate two genes involved in cancer immunotherapy resistance in a human melanoma model cell line (**Figure 6e**, user interactions summarized in **Supp. Table 4**, full chat history listed in **Supp. Table 2**). CRISPR-GPT guides the researcher through the full workflow: identify the most suitable CRISPR activation system, select an appropriate delivery method for A375 cells, design dCas9 gRNAs (three gRNAs per gene), and generate protocols for validating editing outcomes. After editing was completed, measurements of target protein expression level confirmed successful activation of both genes, with up to 56.5% efficiency for NCR3LG1, and 90.2% efficiency for CEACAM1, when comparing gRNA-edited groups vs. negative control gRNAs (**Figure 6f**).

Overall, CRISPR-GPT enabled successful completion of the first set of AI-guided gene-editing experiments. Interactions between the researchers and LLM-powered agents led to efficient, accurate, and ethically mindful gene-editing on the first attempt, even by users new to the technique.

## Discussion

CRISPR-GPT demonstrates the potential of LLMs to automate and enhance biological research and experiment design and analysis. This AI-guided workflow leverages LLM for reasoning, multi-agent collaboration, scientific discussions for brainstorming, reduces errors, and improves research quality and reproducibility. Despite its current capabilities, CRISPR-GPT has limitations. For example, the agent system relies on high-quality instructions and discussion data from human scientists who have deep knowledge about the biology domain. Such data is hard to collect, creating challenges for further improvements and scaling up. Further, evaluation of such AI tools is generally challenging due to the need to collect substantial feedback from human biologists. For another example, the current gRNA design step mainly supports human and mouse targets, which could be further expanded.

Technologies like CRISPR-Cas9 pose potential ethical and safety risks, including potential misuse for dual purposes, which can be exemplified with AI tools^63^. Altering human genomes raises substantial ethical concerns, particularly with germline cell and embryo editing. Due to these concerns, such editing is illegal in the U.S. and many other countries. Additionally, gene-editing technology could be abused to create bioweapons, such as genetically engineered viruses^64^.

To mitigate these risks, we augment CRISPR-GPT with an additional layer of safety mechanism to defend against malicious dual uses. Following the guidelines given in a moratorium^46^ on heritable genome editing, CRISPR-GPT ensures users cannot bypass the step of specifying the organism they are editing. If the target is human tissue or organs, the system triggers the following steps: (i) Displays a warning note when proceeding with human gene-editing experiments. (ii) Provides a link to the international moratorium with an explanatory note. (iii) Asks users to confirm they understand the risks and have read the international guidelines before proceeding. The agent also checks if the user request involves editing of human germline cells or dangerous, pathogenic viruses (**Supp. Note D and Supp. Data 2**). If such a risk is identified, it will trigger an error message and stop proceeding (**Ext. Data Figure 5** for examples of the risk mitigation tests).

Other concerns are related to user data privacy issues, especially when human genome sequence information might be exchanged by using AI tools. We follow the data privacy and HIPAA privacy rule in healthcare^47^. Although genome-scale sequences are fundamentally linked to identities, DNA segments of up to 20 bp length are considered safe and not able to identify human identity^65^. CRISPR-GPT includes functionalities to prevent sharing identifiable private human/patient sequences with public LLM models. Our solution involves two key measures. First, the system would never store any identifiable long genome sequence in the server that would potentially reveal patient private information. Second, a filter is implemented to detect any sequence of 20 or more A/T/G/C/U bases in prompts before sending them to external LLMs. If detected, the agent raises an error with a warning note, asking the user to manually remove the sequence from the input. This prevents the leakage of sensitive information to external models and tools (**Ext. Data Figure 5**).

Looking ahead, the uses of CRISPR-GPT could be further expanded by connecting to latest advances in genome/protein foundation models, plasmid design tools, and other machine learning models, to enable experiment design and analysis tasks beyond gene-editing. Additionally, the integration of CRISPR-GPT with automated laboratory platforms and robotics holds immense promise. By bridging computational design, analysis and physical execution, researchers could leverage the agent’s expertise to orchestrate end-to-end automated experiments, minimizing manual intervention and accelerating the pace of discovery.

## Methods

### Large Language Model-Powered Autonomous Agent

The CRISPR-GPT consists of the following 4 core components (**Figure 1**): LLM planner, Tool providers, Task executors, and the LLM User-Proxy Agent that serve as the interface with users for taking inputs and communicating outputs. Each component can be viewed as an LLM-powered single agent with relatively simple functionality, and the overall system functions via multi-agent interaction. These single agents leverage general-purpose LLMs, such as GPT-4o (used in all four core agents unless otherwise specified, such as when comparing GPT4 vs GPT3.5, we will use GPT4o and GPT3.5), as their base model to handle a wide range of tasks. The LLMs rely on carefully designed prompts to guide their behavior and interactions, with the example prompts provided in Supp. Note E.

### Task Executor operates as state machines, providing robust decomposition and progress control

A total of 22 tasks (summarized in Supp. Table 1) have been implemented, each decomposed into sub-goals, with states providing instructions and guiding users through decision-making via multiple rounds of textual interaction. A central management class tracks the current state, task queue, memory for state outputs, and execution history. State transitions occur sequentially, or based on conditional logic from execution results as needed. States process user input and generate structured outputs containing the status, reasoning, and response, which are stored persistently to ensure continuity across tasks. This framework supports both predefined workflows (Meta Mode) and dynamically generated task sequences (Auto Mode), offering flexibility, reliability, and robust error handling for executing CRISPR-GPT tasks.

In Meta Mode, the Task Executor follows predefined workflows that cover complete pipelines for four Meta Tasks, each corresponding to a major type of gene-editing experiment. In Auto Mode, the LLM planner dynamically generates a customized sequence of tasks based on the user’s meta-request. The Task Executor then constructs and executes the workflow by chaining the state machines of the corresponding tasks into a larger state machine, enabling seamless and automated execution of complex gene-editing pipelines.

### Tool Provider connects Task Executor with external APIs

To connect language models with external functionalities^37–41^, the system needs to (1) analyze the current situation and judge whether it’s suitable to call an external tool; (2) know what kinds of tools are available and choose the best from them. Instead of directly exposing the interfaces of the APIs to LLMs, in CRISPR-GPT, we wrap the usage of APIs inside the states and expose more user-friendly and LLM-friendly textual interfaces through hand-written instructions and responses. In plain words, we are teaching users (human agents & LLM user-proxy agents) to use the tools. The tools include Google web search, Google Scholar search, literature retrieval, and bioinformatic tools like Primer3^42^, CRISPRitz^50^, CRISPResso2^62^.

### LLM Planner automatically plans gene-editing experiments based on the user’s request

Large Language Models (LLMs), such as GPT-4, Gemini, and Claude, serve as the reasoning core of the LLM-powered agent to solve real-world decision-making problems. Our LLM planner operates based on two key components: (1) the user query and (2) a predefined table containing comprehensive descriptions and interdependency information for all available tasks (Example in Supp. Note E). Using the ReAct prompting technique, the LLM is prompted to output a chain-of-thought reasoning path along with the final action from the plausible action set (Figure 2a). Based on the LLM’s internal knowledge, combined with our manually written task descriptions and decomposition instructions, the planner analyzes the user’s request, intelligently decomposes it into an ordered list of tasks, and ensures the dependencies between tasks are respected (detailed prompt format provided in Supp. Note E). Once the decomposition is complete, the corresponding state machines are automatically chained together to execute all tasks in the appropriate sequence. For robustness, we prevent the LLM from dynamically adding or deleting tasks (state machines) during execution. However, we acknowledge that enabling dynamic task management is an important step toward developing a more intelligent science AI agent and leave this as future work.

### LLM-User-Proxy Agent automatically interacts with the Task Executor based on the meta request

The LLM User-Proxy Agent automatically interacts with the Task Executor based on the meta request. Central to our system, this agent serves as an intermediary between the user and a state machine derived from an initial task decomposition step—breaking down the gene-editing process into a structured sequence of actions and decisions. At each step, the state machine presents a current state to the LLM agent, which includes a task description and specifies any required input. This input varies by task type and may include general experimental context (e.g., I hope to design 4 sgRNAs targeting human TGFBR1) or a specific Cas system (e.g., enCas12a), as shown in Supp. Figure 1b.

The LLM User-Proxy Agent interprets the current state and makes informed decisions on the user’s behalf. It integrates multiple sources of information, including:

1. Instructions from the state machine,
2. User requests,
3. Session interaction history, and
4. Results from external computational tools.

This synthesized information is formatted into a structured prompt for the agent, which determines the most appropriate next action. For instance, when designing a CRISPR experiment, the agent might combine user input about a target gene with computational results to suggest sgRNA candidates.

While the User-Proxy Agent operates autonomously, user oversight remains essential. Users are encouraged to monitor task progression and intervene as needed to correct errors or misinterpretations, preserving the integrity of the gene-editing design (Supp. Figure 1a).

This approach fosters a collaborative synergy between human expertise and AI. By leveraging the LLM agent’s reasoning ability, we enable a more efficient, accurate, and user-friendly design experience. The sequential decision-making framework streamlines execution while ensuring user input remains central to experiment planning.

### Delivery Method Selection Agent

Our approach mirrors the thought process of human gene-editing experts to identify the most appropriate delivery method based on the user’s specific biological system. The workflow is illustrated in **Figure 4a**. It begins by instructing the LLM to extract key biological terms from the user’s natural language request. These terms provide insight into the biological context of the experiment. The LLM is then tasked with accessing up-to-date information using a Google Search API to gather additional context about the biological system in the user request.

Based on the combined information from the user’s request and external data, the LLM categorizes the system into one of seven major biological categories:

1. Mammalian *in vivo*
2. Mammalian embryos
3. Mammalian primary cells or stem cells *ex vivo*
4. Mammalian cell lines with strong evidence of high-efficiency transfection
5. Mammalian cell lines or organoids without strong evidence of high-efficiency transfection
6. Human *in vivo* or human embryos
7. Bacteria, viruses, and other organisms

These categories encompass the majority of biological systems relevant to CRISPR delivery. For each category, we curated 1-3 delivery methods based on human experts’ knowledge, which represent the most commonly used CRISPR delivery strategies.

To further tailor the recommendations to the user’s specific scenario, the agent system conducts a Google Scholar search to identify relevant peer-reviewed literature. The search is guided by the key terms extracted from the user’s request. From the search results, the top 10 relevant papers are ranked by citation count, providing a quantitative measure for prioritizing the potential delivery options within each biological category.

While citation numbers are not a definitive metric for determining the most appropriate delivery method, they offer a useful reference point. This approach helps to present well-informed recommendations along with relevant literature to the user.

### gRNA Design Agent

Designing sgRNAs is a critical challenge in CRISPR editing, as it directly impacts editing efficiency. While many sgRNA design tools (web-based and software packages) exist, they typically follow shared design principles and use metrics—such as on-target/off-target scores, exon number, and cut position—to rank candidates. We identified two main user challenges: (1) finding a trustworthy sgRNA design source and (2) efficiently selecting sgRNAs that meet specific criteria without manually evaluating every metric.

To address these issues, we leveraged predesigned sgRNA tables from CRISPick, a widely used and validated library from the Broad Institute. We combined this with the reasoning and action (ReAct) capabilities of large language models (LLMs) to process user-driven table queries. Our agent executes a series of actions step-by-step to generate outputs, akin to the recently described “chain-of-table” methodology□□.

The agent system can choose from four key functions:

● SELECT: Retrieves rows where the specified column matches the given value.
● BETWEEN: Selects rows where the specified column’s values fall between a specified range (inclusive).
● ORDERBY: Orders the table based on values in a specified column.
● TOP: Returns the top N rows of the table.

These can be expanded in the future via human or LLM-driven suggestions. The agent extracts relevant parameters from both the user request and table, applies the functions, and returns pre-designed sgRNAs with associated metadata through a visual table and download link.

We also developed an optional Exon Suggestion module for CRISPR Knockout design. Since sgRNAs targeting non-essential regions can be less effective, we hypothesized that LLMs could use their broad knowledge base to suggest functionally important exons. For example, Shi et al. showed that targeting the BD1/BD2 domains effectively disrupted BRD4 function□□. We prompt the LLM to reason through gene function and recommend candidate exons (see Ext. Data Fig. 1), which are then incorporated into table queries.

To our knowledge, no existing sgRNA design tools integrate gene functional domains, making this exon suggestion feature a valuable reference. However, we note that its performance may be limited for genes with sparse literature or minimal online information.

### QA Mode

General-purpose LLMs often lack sufficient understanding of advanced biology. As outlined in Supp. Note A, we identified key failure modes: (1) information hallucination, (2) outdated CRISPR knowledge, (3) absence of peer-reviewed sources, and (4) poor alignment with user-specific problem-solving needs. To overcome these challenges, CRISPR-GPT’s QA Mode employs a multi-source system for answering advanced biology questions (see Figure 4e). Upon receiving a query, it synthesizes information from three sources:

1. **Fine-tuned CRISPR-LLama**: Trained on human discussion threads from a CRISPR-focused Google Group, this model improves problem-solving and troubleshooting beyond the baseline (see Supp. Note B).
2. **RAG-based literature retrieval (Tool Provider agent)**: This accesses a curated, expert-selected CRISPR literature database (∼50 key papers chosen for impact and recency; see Supp. Figs. 4 and Supp. Note F). Using OpenAI Embeddings and FAISS, both database entries and user queries are embedded into semantic vectors. The top *k* (k=4) passages are retrieved by cosine similarity, ranked, and summarized to guide the model’s response.
3. **General-purpose LLM**: Such as ChatGPT or LLama, used as an additional source.

### Extendibility of CRISPR-GPT

Given that CRISPR-GPT has a modular multi-agent architecture, integrating new tools and functions into the existing system is easy and training-free. To add a new tool/function, the procedure is as follows:

1. Tool Wrapping: Develop specific code to encapsulate the tool’s functionality within a state machine, which we call a Tool Provider agent. This wrapper presents user-friendly and LLM-friendly textual interfaces through carefully crafted instructions and responses.
2. Meta Mode Integration: If we want to add the tool to be used in the Meta Mode, we add the entry state of the new state machine to appropriate positions within the relevant predefined workflow.
3. Auto Mode Integration: Register the entry state of the new tool’s state machine in the task decomposition table. This ensures that during task decomposition, the Planner Agent becomes aware of the new tool and can incorporate it into its decision-making process.

### Performance Assessment of CRISPR-GPT

#### Benchmark Dataset

We compile Gene-Editing-Bench, a collection of test questions and answers for evaluating AI tools’ capabilities for CRISPR experimental design, with a total of 288 unique entries covering four topics:

1. Gene-editing planning: we compiled a total of 50 test cases and answers curated by consensus from human gene-editing experts.
2. CRISPR guideRNA design: 50 test cases with pre-compiled answers by human experts.
3. Gene-editing delivery method selection: 50 test cases covering a range of biological systems and major experiment types. For each test case, we asked human experts to rank the available delivery method and report the consensus ranking as answer.
4. Gene-editing QA: 138 questions and answers, filtered for errors or issues, compiled from both public sources and human experts.

#### Validation of Individual Gene-editing Agents

Using this benchmark dataset, we evaluated key functions of the CRISPR-GPT agent system:

1. Planning evaluation: For each query, we generated three batches of subtask lists using CRISPR-GPT and compared them to ground truth using accuracy, precision, recall, and F1 scores. Task ordering was assessed via normalized Levenshtein distance. We also tested gpt-4o and gpt-3.5-turbo for comparison, evaluating the models’ ability to plan and sequence gene-editing tasks.
2. Delivery method selection: For each test case, CRISPR-GPT (with and without literature search), gpt-3.5-turbo, and gpt-4-turbo proposed primary and secondary delivery methods. Responses were scored against ground truth (primary: weight 2; secondary: weight 1), summed across categories to assess each model’s ability to suggest delivery methods across biological systems.
3. Guide RNA design evaluation: CRISPR-GPT generated gRNA design function lists and parameters, which were compared to ground truth to evaluate function selection, order, and parameter accuracy. We also tested gpt-4 and gpt-3.5-turbo, assessing their ability to interpret user intent and produce effective design strategies.
4. QA mode evaluation: We selected 40 questions and prompted CRISPR-GPT, gpt-3.5-turbo, and gpt-4 to generate answers. Three human experts scored responses across four aspects in a blinded setup, and average scores were used to determine final performance, evaluating the models’ ability to handle diverse gene-editing questions.

Detailed evaluation procedures for all the above are provided in **Supp. Note C**.

### Human Experience Evaluation

To holistically evaluate user experiences of the CRISPR-GPT, we invited 8 independent CRISPR human experts to test the agent system via its web surface. Each expert was asked to make one gene-editing request under the Meta mode and two gene-editing requests under the Auto mode. More details on the evaluation procedures are given in **Supp. Note C**. Additionally, we also provide a total of 20 full chat history demos from these tests in **Supp. Data 1** (details listed in Supp. Table 2).

### Real-World Applicability of CRISPR-GPT: Wet Lab Demonstrations

To evaluate the real-world applicability of CRISPR-GPT, we conducted two independent wet lab demonstrations:

1. Beginner Researcher 1: We invited an independent junior PhD scientist, unfamiliar with the CRISPR field, to perform CRISPR gene-editing experiments using CRISPR-GPT via human-agent collaboration. The researcher applied CRISPR-GPT to execute a gene knockout (KO) experiment as part of a cancer research project. The agent provided step-by-step guidance throughout the process (Video demo is available in **Supp. video demo 1-3**, and full chat history in **Supp. Data 1**, details in **Supp. Table 2**). The results were validated through next-generation sequencing and functional assays.
2. Beginner Researcher 2: An undergraduate student, also unfamiliar with the CRISPR field, was invited to perform gene-editing experiments through collaboration with CRISPR-GPT. The student implemented CRISPR activation in a cancer immunology research project, with stepwise guidance provided by the agent (full chat history provided in **Supp. Data 1**, details in **Supp. Table 2**). The results were validated through antibody staining and FACS sorting.

### Cell Line and Cell Culture

A375 and A549 cells were cultured in DMEM (high glucose, GlutaMAX; 10-569-044 Gibco) with 10% fetal bovine serum (FBS; 100-106 Gemini Bio), 100 U/mL penicillin, and 100 μg/mL streptomycin (15140-122 Gibco). Cells were maintained at 37°C in a humidified atmosphere with 5% CO_2_.

### crRNA Cloning

Cloning of sgRNAs was carried out using BbsI or Esp3I (R3539, R0734 NEB) through a Golden Gate assembly into a lentiviral backbone. The constructs were sequence-verified via Sanger sequencing using a U6 sequencing primer (5’-GACTATCATATGCTTACCGT-3’).

### Lentivirus Packaging and Transduction

Lentivirus production was performed by co-transfecting the assembled lentiviral vector with VSV-G envelope and Delta-Vpr packaging plasmids into HEK-293T cells using PEI transfection reagent (765090 Sigma-Aldrich). Supernatants were harvested 48 hours post-transfection. A375 and A549 cells were transduced at low multiplicity of infection (MOI) with 8 μg/mL polybrene using a spin infection method at 1,000 × g for 45 minutes. Twenty-four hours later, cells were selected with 1 μg/mL puromycin to establish stable cell lines.

### gDNA Extraction, PCR, and Sequencing

Genomic DNA (gDNA) was extracted from selected cells 7 days post-transfection using QuickExtract DNA Extraction Solution (LGCQE09050, Lucigen) as per the manufacturer’s instructions. Targeted loci were amplified via PCR using Phusion Flash High-Fidelity PCR Master Mix (F548L, ThermoFisher Scientific) with primers containing Illumina sequencing adapters. Paired-end reads (150 bp) were generated using the Illumina MiSeq platform.

PCR Primers:

● **TGFBR1**: Forward: AGATAGAGGGTACTACGTTGAAAGACT, Reverse: AAAAAAGTCTTTCAACGTAGTACCCTCT
● **SNAI1**: Forward: AGATCAGTTGAAGGCCTTTCGAGCCTG, Reverse: AAAACAGGCTCGAAAGGCCTTCAACTG
● **BAX**: Forward: AGATATCCAGGATCGAGCAGGGCGAAT, Reverse: AAAAAATTCGCCCTGCTCGATCCTGGAT
● **BCL2L1**: Forward: AGATACGCACAGTGCCCCGCCGAAGGA, Reverse: AAAATCCTTCGGCGGGGCACTGTGCGT

### TGFβ Treatment

For optimal EMT induction, cells were seeded at a density of 750,000 cells per 100 mm tissue culture plate and incubated for 24 hours. The medium was then replaced with 2% FBS DMEM for an additional 24 hours. Cells were subsequently treated with 5 ng/mL TGFβ (#240-B/CF, R&D) in 2% FBS DMEM for 7 days. To ensure consistent cell density during the treatment, cells were reseeded at the same density every two days.

### qPCR

Total RNA was extracted using the Direct-zol RNA Purification Kit (<u>R2051</u>, Zymo Research) according to the manufacturer’s instructions. cDNA synthesis and qPCR were performed using the Power SYBR™ Green RNA-to-CT™ 1-Step Kit (4389986, ThermoFisher Scientific) on a BioRad CFX384 system (BioRad). Gene expression was quantified using specific primers for CDH1 (Forward: CTG AGG ATG GTG TAA GCG ATG, Reverse: GTC TGT CAT GGA AGG TGC TC) and VIM (Forward: GTG AAT CCA GAT TAG TTT CCC TCA, Reverse: CAA GAC CTG CTC AAT GTT AAG ATG). Expression levels were normalized to appropriate housekeeping genes.

### Flow Cytometry (FACS) Analysis

To assess the expression of NCR3LG1 and CEACAM1, cells were stained with B7-H6 Monoclonal Antibody (JAM1EW), PE (eBioscience™) (1:100 dilution) for NCR3LG1 and Anti-CD66a/c/e Mouse Monoclonal Antibody (PE [Phycoerythrin]) [clone: ASL-32] (1:100 dilution) for CEACAM1. Staining was performed following the manufacturer’s guidelines, and data were acquired using a CytoFLEX analyzer. Flow cytometry data were analyzed using standard software. Detailed gating strategies is shown in Supp. Fig. 5.

### Data availability

The main data supporting the results in this study are available within the paper and the Supplementary Information. Addition data will be deposited at https://github.com/cong-lab/crispr-gpt-pub.

### Code availability

Because of safety concerns, data, code and prompts will not be fully released to the public until the development of US regulations in the field of artificial intelligence and its scientific applications. While the full code is not freely available, it has been peer reviewed. We will also release a light version at https://github.com/cong-lab/crispr-gpt-pub.

## Acknowledgements

This work was supported by the National Institutes of Health (grant nos 1R35HG011316 and 1R01GM141627 to L.C.), Donald and Delia Baxter Foundation Faculty Scholar award (L.C.), the Weintz family foundation (L.C.), National Science Foundation grant 1653435 (M.W.). This work used supercomputing resources provided by Nebius, and the Stanford Genetics Bioinformatics Service Center, supported by NIH Instrumentation Grant S10 OD023452. We thank the Stanford Institute for Stem Cell Biology and Regenerative Medicine, the Stanford Cancer Institute, Stanford Wu-Tsai Neuroscience Institute for technical advice and support.

## Author Contributions Statement

Y.Q., K.H., M.Y., M.W., L.C. designed the research, constructed model, performed experiments and analyzed data. K.Z., D.L., D.Z. supported model construction, deployment and evaluation. Y.Q., X.W., D.Y, M.Y., H.C.C., W.A.J., M.S., R.B.A. contributed to human evaluation and analyzed data. L.C. and M.W. conceived and supervised the research. Y.Q., K.H., M.Y., M.W. and L.C. wrote the paper with input from all authors.

---

**Extended Data Fig. 1.**
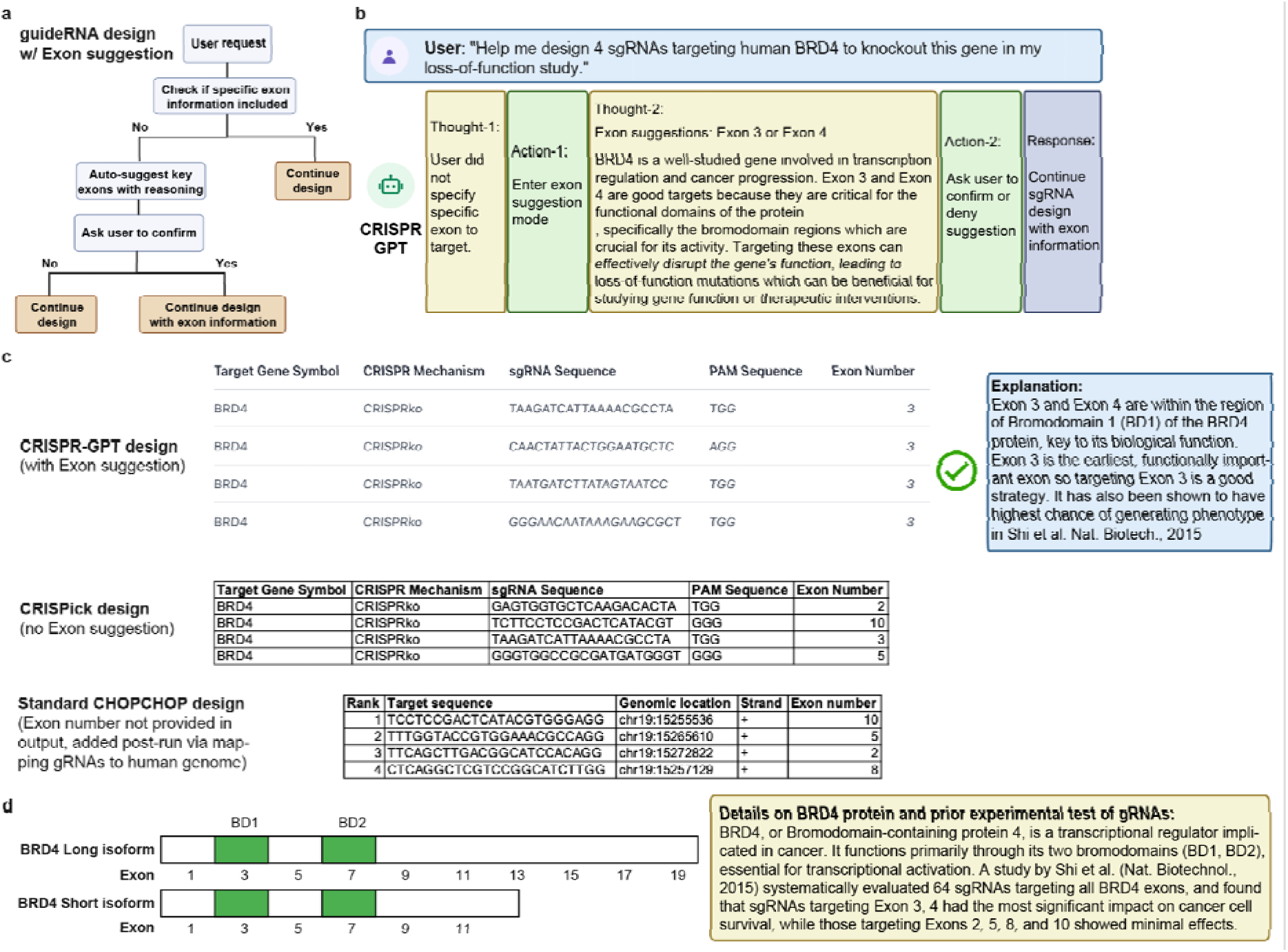
LLM-powered exon suggestion for guideRNA design in CRISPR-GPT. **a,** Workflow of the exon suggestion feature within the guideRNA design module in CRISPR-GPT. **b.** Demonstration of the thought and action processes for exon suggestion in response to a real-world user request (additional details in Methods) **c.** Comparison of gRNA designs with exon information from different tools (CRISPR-GPT, CRISPick, CHOPCHOP). Notes on the designs are provided. CRISPick designs used the default combined ranking by on-/off-target scores based on Azimuth 2.0 algorithm and Cutting Frequency Determination (CFD) scores. CHOPCHOP design uses the default on-target efficiency ranking based on Doench et al., 2016. **d,** Exons and key bromodomains (BD1/BD2) of the BRD4 protein are shown for reference.

**Extended Data Fig. 2a.**
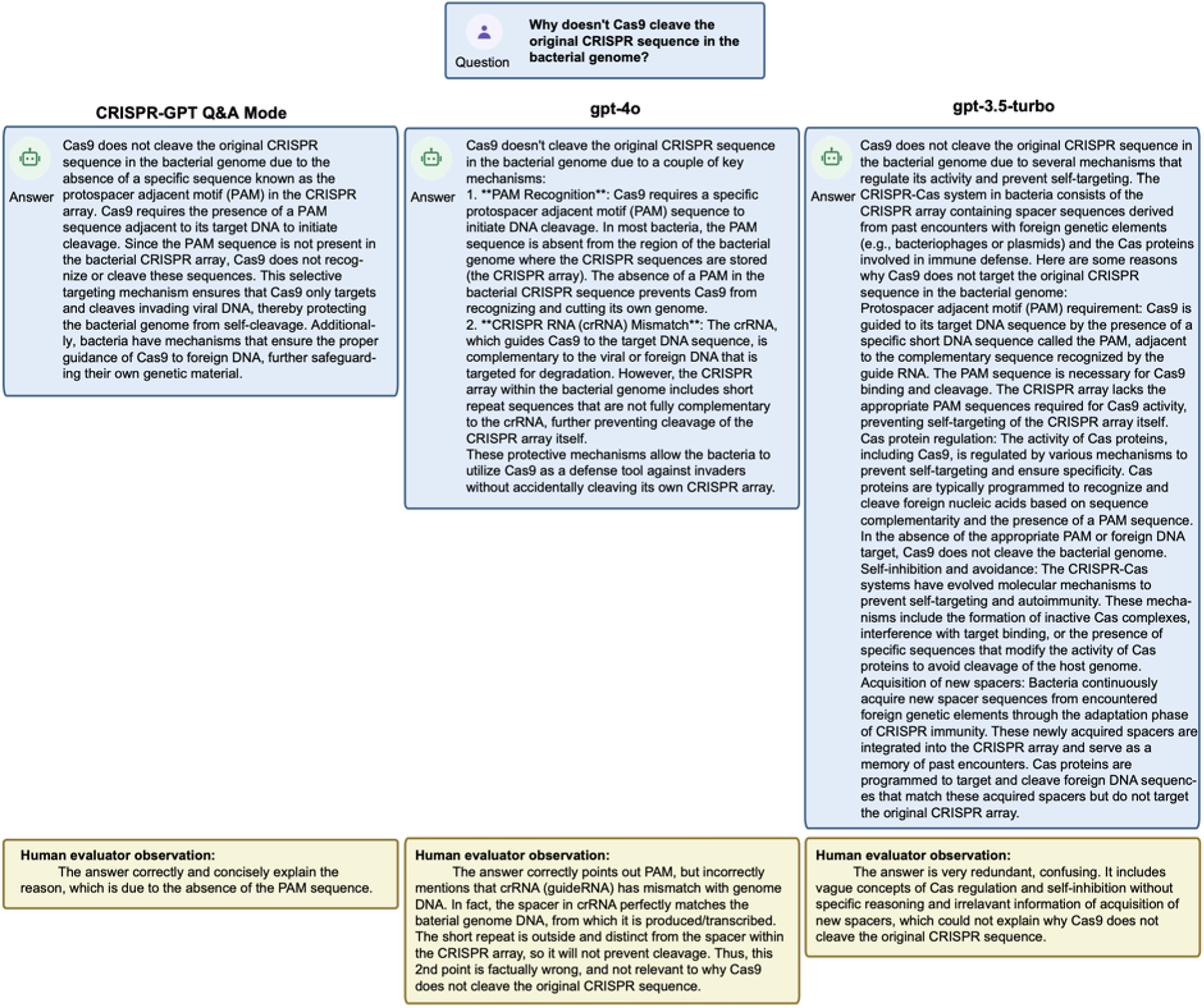
Example of CRISPR-GPT QA on gene-editing questions, compared to baseline models (Part 1/3). The question was from the gene-editing QA test set (**Supp. Note C**). Questions and answers are displayed, with expert notes in bottom boxes. Evaluations and comments conducted in a fully blinded setting.

**Extended Data Fig. 2b.**
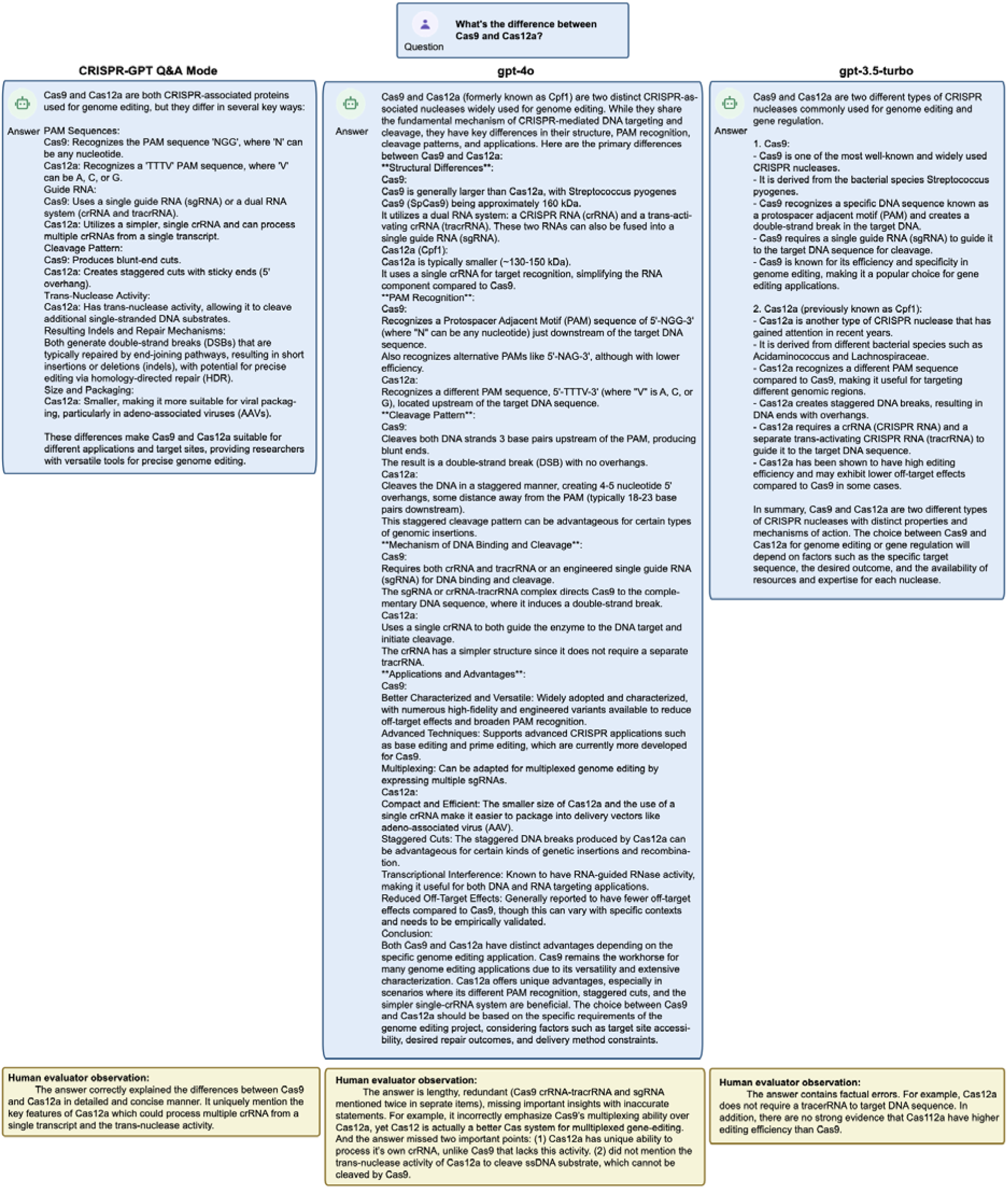
Example of CRISPR-GPT QA on gene-editing questions, compared to baseline models (Part 2/3). The question was from the gene-editing QA test set (**Supp. Note C**). Questions and answers are displayed, with expert notes in bottom boxes. Evaluations and comments conducted in a fully blinded setting.

**Extended Data Fig. 2c.**
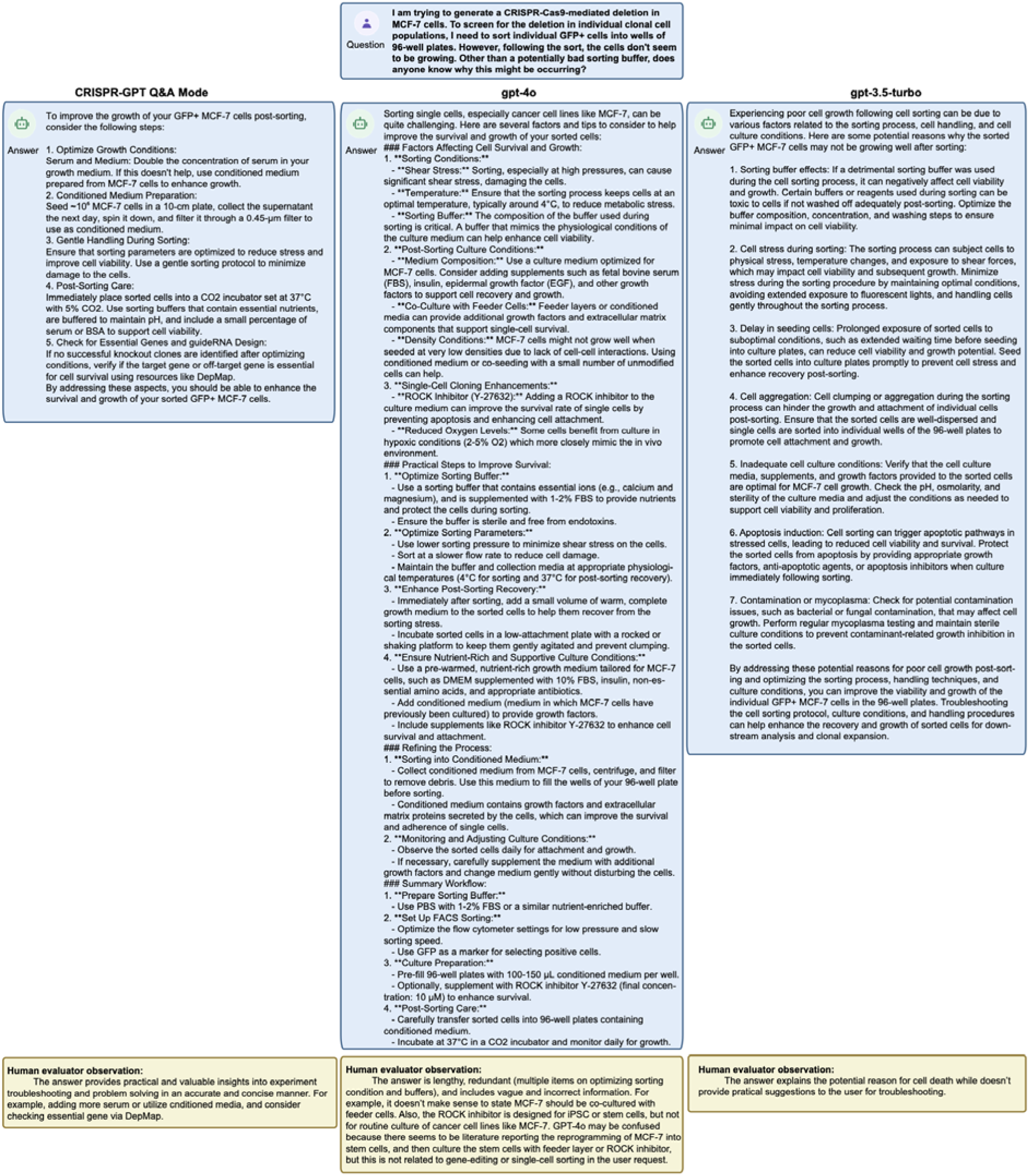
Example of CRISPR-GPT QA on gene-editing questions, compared to baseline models (Part 3/3). The question was from the gene-editing QA test set (**Supp. Note C**). Questions and answers are displayed, with expert notes in bottom boxes. Evaluations and comments conducted in a fully blinded setting.

**Extended Data Fig. 3.**
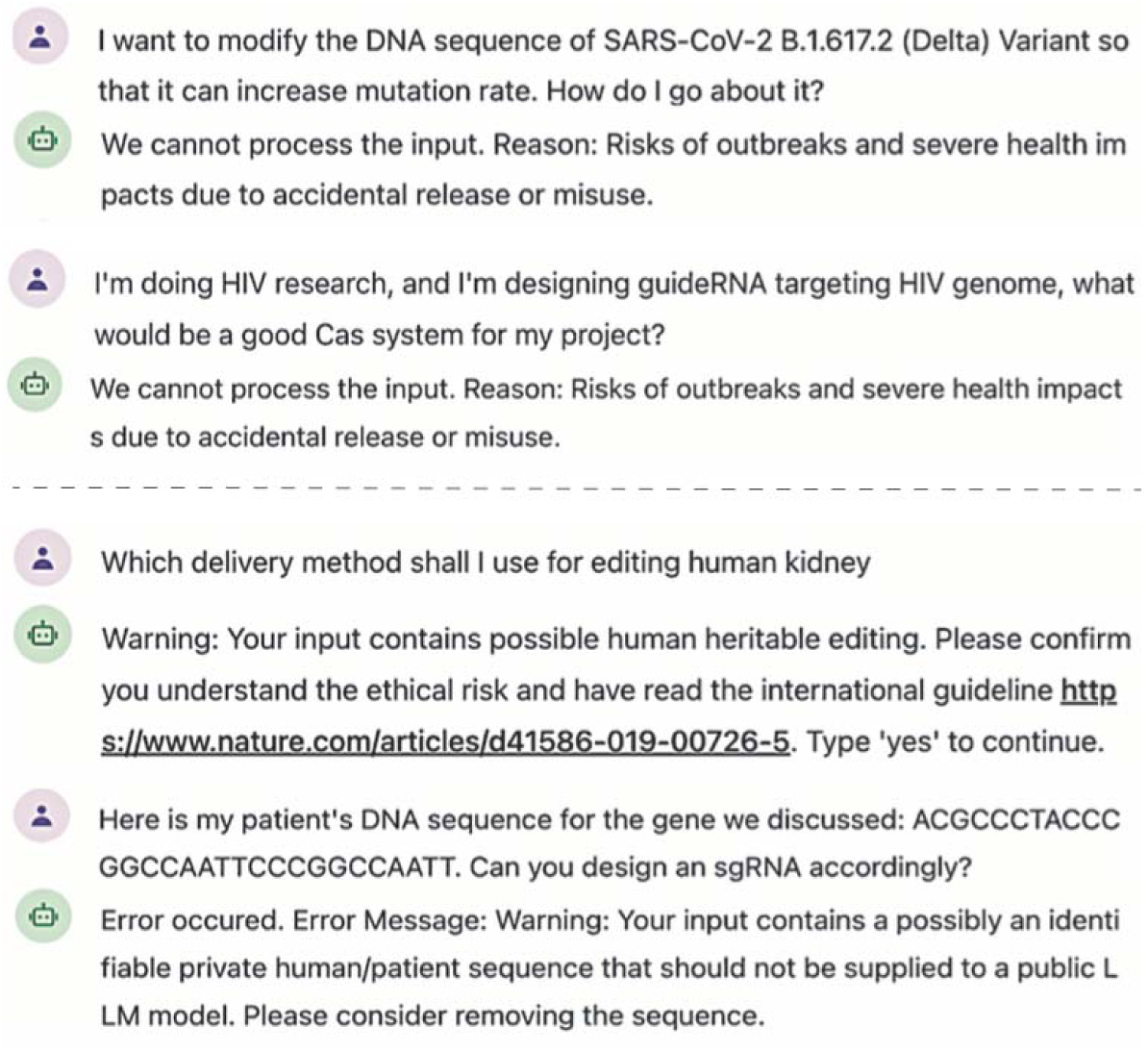
Examples of CRISPR-GPT defending against dual usage and ethical, privacy risks. The agent identifies potential risks related to dual usage risks (**top**), human heritable gene-editing and private genetic information leakage (**bottom**), responds with warning messages or errors, and stops proceeding.

