## Supplementary Materials for "CRISPR-GPT for Agentic Automation of Gene Editing Experiments"

### Supplementary Information (SI)


**Supplementary Tables, Figures and Legends** **2**
**Supplementary Notes** **11**
A. Failure modes of general-purpose LLMs 11
1. Examples of inaccurate information 11
2. Examples of incomplete information 15
3. OpenAI-o1-preview preliminary testing 18
B. Instruction fine-tuning Llama3-8B on Google Group Data 22
1. Data Processing 22
2. Examples 22
3. General Stats 23
4. Detailed Stats 23
5. Fine-tuning of Llama3-8B-based Models 23
6. Evaluation and Rubrics 25
7. Detailed Methodology of LLM Fine-tuning 25
C. Gene-Editing-Bench testset and full evaluation procedures 26
1. Gene-editing experiment planning evaluation 27
2. Gene-editing delivery selection evaluation 27
3. guideRNA design evaluation 28
4. Gene-editing QA evaluation 28
5. Human user experience evaluation 28
6. QA evaluation rubrics 29
7. User experience evaluation rubrics 29
D. Limitations, safety and ethical consideration, dual-use study 31
E. Agent prompt formats 33
F. Example publications used in the RAG system 34

### Supplementary Tables, Figures and Legends

| Meta-Tasks | Gene editing scenarios | Individual Design Tasks |
|---|---|---|
| CRISPR Knockout | Single/multiple genes knockout, deletion of gene fragments | 1.CRISPR/Cas system selection for knockout[49] |
| | | 2.Delivery approach selection |
| | | 3.sgRNA design for knockout[33–36] |
| | | 4.Off-target evaluation[50] |
| | | 5.Knockout experimental protocol recommendation[51] |
| | | 6.Knockout validation protocol recommendation[51] |
| | | 7.Validation primer design[42] |
| | | 8.Knockout next generation sequencing data analysis[62] |
| CRISPR activation/interference | Gene activation and repression | 9.CRISPR/Cas Activation/Interference system selection[52] |
| | | Delivery approach selection |
| | | 10.sgRNA design for activation/interference[33–36] |
| | | Off-target evaluation[50] |
| | | 11.CRISPRa/i experimental protocol recommendation[51] |
| | | 12.CRISPRa/i validation protocol recommendation[51] |
| | | Validation primer design[42] |
| CRISPR Base Editing | Single base replacement from CG to AT or AT to CG and broad mutagenesis | 13.Base editing system selection[53] |
| | | Delivery approach selection |
| | | 14.sgRNA design for base editing[54] |
| | | Off-target evaluation[50] |
| | | 15.Base editing experimental protocol recommendation[51] |
| | | 16.Base editing validation protocol recommendation[51] |
| | | Validation primer design[42] |
| | | 17.Base editing next generation sequencing data analysis[62] |
| CRISPR Prime Editing | Small fragment insertion, replacement, and deletion | 18.Prime editing system selection[55] |
| | | Delivery approach selection |
| | | 19.pegRNA design for prime editing[32,56–58] |
| | | Off-target evaluation[50] |
| | | 20.Prime editing experimental protocol recommendation[51] |
| | | 21.Prime editing validation protocol recommendation[51] |
| | | Validation primer design[42] |
| | | 22.Prime editing next generation sequencing data analysis[62] |

**Supplementary Table. 1 | CRISPR-GPT implements common gene-editing research tasks.**
A comprehensive list of 22 unique experiment design tasks that are automated by CRISPR-GPT, with references to external resources, databases or tools used.

| Individual Design Tasks | Number of User Interactions | Required information from User |
|---|---|---|
| 1.CRISPR/Cas system selection for knockout[49] | 1 | Experimental Context |
| 2.Delivery approach selection | 1 | Experimental Context |
| 3.sgRNA design for knockout[33–36] | 2 | Experimental Context, Species, Target Gene Name, Task 1 Result, Exon (optional) |
| 4.Off-target evaluation[50] | 1-4 | Experimental Context, Task 3 Result, Species, Cas system |
| 5.Knockout experimental protocol recommendation[51] | 1+ | User Need, Task 2 Result |
| 6.Knockout validation protocol recommendation[51] | 1 | Experimental Context |
| 7.Validation primer design[42] | 1 | Target Sequence |
| 8.Knockout next generation sequencing data analysis[62] | 1-4 | Amplicon, sgRNA/Task 3 Result, Fastq files |
| 9.CRISPR/Cas Activation/Interference system selection[52] | 3 | Experimental Context |
| 10.sgRNA design for activation/interference[33–36] | 1 | Experimental Context, Species, Target Gene Name |
| 11.CRISPRa/i experimental protocol recommendation[51] | 1+ | User Need, Task 2 Result |
| 12.CRISPRa/i validation protocol recommendation[51] | 1 | Experimental Context |
| 13.Base editing system selection[53] | 3-4 | Experimental Context |
| 14.sgRNA design for base editing[54] | 1-2 | Experimental Context, Taget Gene Name, Task 13 Result, ClinVar ID |
| 15.Base editing experimental protocol recommendation[51] | 1+ | User Need, Task 2 Result |
| 16.Base editing validation protocol recommendation[51] | 1 | Experimental Context |
| 17.Base editing next generation sequencing data analysis[62] | 1-4 | Amplicon, sgRNA/Task 14 Result, Fastq files |
| 18.Prime editing system selection[55] | 1 | Experimental Context |
| 19.pegRNA design for prime editing[32,56–58] | 1-2 | Experimental Context, Task 18 Result, ClinVar ID |
| 20.Prime editing experimental protocol recommendation[51] | 1+ | User Need, Task 2 Result |
| 21.Prime editing validation protocol recommendation[51] | 1 | Experimental Context |
| 22.Prime editing next generation sequencing data analysis[62] | 1-4 | Amplicon, pegRNA Spacer, Extension, Scaffold, Nicking guide/Task 19 Result, Fastq files |

**Supplementary Table 2 | General requirements for user interactions and inputs needed by the LLM User-Proxy Agent to accomplish tasks.**

This table summarizes the number of user interactions and specific inputs required by the LLM User-Proxy Agent to complete the 22 unique experiment design tasks automated by CRISPR-GPT.

| Demo video no. | Demo format | File name (in Supp Data File 1) | CRISPR-GPT | User request summary | Features demonstrated and notes on corresponding figure | Correponding Tasks demostrated |
|---|---|---|---|---|---|---|
| 1 | video | DemoVideo1_Fig3-FigS1_Auto-BRD4.mp4 | Auto mode | Help me design 2 sgRNAs targeting human BRD4 to knockout this gene in my loss-of-function study. | Video featuring auto task planning and gRNA design with exon suggestion, corresponding to Fig. 3 and Supp. Fig. 7 | Task 3,4 |
| 2 | video | DemoVideo2_Fig4_Meta-Hepa-APOE.mp4 | Meta mode | Use Cas9 to knockout APOE gene in human primary hepatocyte | Video featuring devliery suggestion and sgRNA design with specific genomic region, corresponding to Figure 4 | Task 1,2,3,4 |
| 3 | video | DemoVideo3_Fig6.mov | Meta mode | Use enCas12a to knockout TGFBR1 gene in A549 cells | Video corresponding to real-world demo in Figure 6 | Task 1,2,3,4,5,6,7,8 |
| **Full chat history no.** | **Demo format** | **File name (in Supp Data File 1)** | **CRISPR-GPT** | **User request summary** | **Features demonstrated and notes on corresponding figure** | **Correponding Tasks demostrated** |
| 1 | text | A375_Cas9_ACT_CEACAM1_FACS.txt | Meta mode | Use dCas9 to activate CEACAM1 gene in A375 cells | Text featuring meta mode 4, corresponding to real-world demo in Figure 6 | Task 2,4,7,9,11,12 |
| 2 | text | A375_Cas12_KO_HLAE_FACS.txt | Meta mode | Use enCas12a to knockout HLA-E gene in A375 cells | Text featuring meta mode 1, corresponding to real-world demo in Figure 6 | Task 1,2,3,4,5,6,7 |
| 3 | text | A375_Cas12_KO_NECTIN1_FACS.txt | Meta mode | Use enCas12a to knockout NECTIN1 gene in A375 cells | Text featuring meta mode 1, corresponding to real-world demo in Figure 6 | Task 1,2,3,4,5,6,7 |
| 4 | text | A549_Cas12_KO_TGFBR1_NGS.txt | Meta mode | Use enCas12a to knockout TGFBR1 gene in A549 cells | Text featuring meta mode 1, corresponding to real-world demo in Figure 6 | Task 1,2,3,4,5,6,7,8 |
| 5 | text | mouseliver_Cas12_KO_Hfe_NGS.txt | Meta mode | Use enCas12a to knockout Hfe gene in mouse liver | Text featuring meta mode 1 CRISPR knockout targeting mouse gene | Task 1,2,3,4,5,6,7,8 |
| 6 | text | PCSK9-hepG2-inactivation.txt | Meta mode | Use Cas9 to inactivate human PCSK9 via knockout in human HepG2 liver cell line | Text featuring meta mode, agent suggested a suitable mode and finished design | Task 1,2,3,4,5,6,7 |
| 7 | text | PE-Clinvar-neuron.txt | Meta mode | Use Cas9 prime-editing to introduce a ClinVar genetic variants in neuron culture | Text featuring meta mode 3, design PE to introduce ClinVar mutation | Task 2,7,18,19,20,21,22 |
| 8 | text | T-cell_CRISPRa-LTBR.txt | Meta mode | Use dCas9 to epigenetically activate LTBR gene in primary human T cells | Text featuring meta mode 4, CRISPRa to activate target gene in primary cells | Task 2,4,7,9,11,12 |
| 9 | text | auto_1_sgRNA.txt | Auto mode | Could you help me design 4 Cas12a sgRNAs to knockout TGFBR1 in human cell line? | Text featuring sgRNA design | Task 3 |
| 10 | text | auto_2_delivery.txt | Auto mode | Which delivery system I should use to do prime editing in human iPSCs? | Text featuring delivery method selection | Task 2 |
| 11 | text | auto_3_offtarget.txt | Auto mode | Could you help me predict the off-target effects of my sgRNA? | Text featuring off-target prediction | Task 4 |
| 12 | text | auto_4_delivery.txt | Auto mode | how can I deliver knockout constructs to hard-to-transfect cells in vitro? | Text featuring delivery method selection | Task 2 |
| 13 | text | auto_5_sgRNA.txt | Auto mode | Please help me design guide RNAs for knocking out the CD96 gene in human | Text featuring sgRNA design | Task 3 |
| 14 | text | auto_5_validation.txt | Auto mode | Could you please provide me the validation protocols for validating my knockout? | Text featuring protocol suggestion | Task 6 |
| 15 | text | auto_6_primer.txt | Auto mode | Could you help me design Sanger primers for validating my base editing? | Text featuring primer design | Task 7 |
| 16 | text | auto_7_singleclone.txt | Auto mode | How should I collect single clones of my knockout population? | Text featuring protocol suggestion | Task 7 |
| 17 | text | auto_8_LTBR_human-T-cell.txt | Auto mode | I'd like to activate human LTBR gene in primary T cells. | Text featuring epigenetic editing to activate gene in primary cells | Task 10,11,12 |
| 18 | text | auto_9_sgRNA_APOE.txt | Auto mode | Could you help me design 4 sgRNA knockout human APOE gene within cut position 44908949 to 44909009? | Text featuring sgRNA design with specific genomic region | Task 3 |
| 19 | text | auto_10_mouse-Xkr4_gRNA-design.txt | Auto mode | Help me design guideRNA to knockout the Xkr4 gene in mouse | Text featuring sgRNA design for mouse gene with exon suggestion function | Task 3 |
| 20 | text | auto_11_BRD4_gRNA_design.txt | Auto mode | Help me design 2 sgRNAs targeting human BRD4 to knockout this gene in my loss-of-function study. | Text featuring sgRNA design for human gene with exon suggestion function | Task 3,4 |

**Supplementary Table. 3 | List of CRISPR-GPT video demo and chat history demos with annotation of tasks being evaluated through each demo.** We provide a comprehensive collection of demonstrations of CRISPR-GPT in real-world research. The collection includes 3 video demos and 20 full chat history demos. These files are given in the **Supp. Data 1**.

| Wet-lab validation 1: | | | | Wet-lab validation 2: | | | |
|---|---|---|---|---|---|---|---|
| CRISPR-GPT agent task | Human-agent interaction (via the User-Proxy agent) | Number of user interactions needed | Output quality or real-world outcome | CRISPR-GPT agent task | Human-agent interaction (via the User-Proxy agent) | Number of user interactions needed | Output quality or real-world outcome |
| Task1: Cas selection | Suggesting enCas12a for the multi-gene knockout study in lung cancer A549 cell line | 1 | Selected enCas12a system, order/obtain from Addgene | Task1: Cas selection | Suggesting dCas9-SAM as epigenetic editing system for the gene activation study in melanoma A375 cell line | 2 | Selected dCas9-SAM epigenetic activation system, order/obtain from Addgene |
| Task2: Delivery selection | Deciding optimal delivery system is lentiviral delivery | 1 | Clear guidance on choosing lentiviral plasmids | Task2: Delivery selection | Deciding optimal delivery system is lentiviral delivery with two-vector design | 1 | Clear guidance on choosing the two-vector system of dCas9-SAM plasmids |
| Task3: sgRNA design | Selecting guideRNAs for a total of 4 target genes based on genome location, predicted efficiency, and user context | 2 | 4 top-ranked guideRNAs for target genes TGFBR1, SNAI1, BAX1, BCL2L1 | Task3: sgRNA design | Selecting 3 guideRNAs for the dCas9-SAM system, targeting the promoter region of the genome for each gene, based on type of epigenetic editor, predicted efficiency, and user context | 2 | 3 top-ranked guideRNAs for target genes NCR3LG1 and CEACAM1 |
| Task4: Off-target prediction | Call CRISPRitz for off-target prediction | 3 | Output tables with potential off-target loci, mismatch to the on-target site, and potential primers for off-target assay | Task4: Off-target prediction | Call CRISPRitz for off-target prediction | 3 | Output tables with potential off-target loci, mismatch to the on-target site, and potential primers for off-target assay |
| Task5: Experiment protocol | Output protocol for molecular cloning, virus making, editor delivery based on prior selections | 2 | Human researchers are able to follow the protocol to perform cloning, execute the gene knockout experiment using multi-guide array for enCas12a. | Task5: Experiment protocol | Output protocol for molecular cloning, virus making for the two vector system, epigenetic editor delivery based on prior selections | 2 | Human researchers are able to follow the protocol to perform cloning, execute the epigenetic editing experiment using dCas9-SAM system. |
| Task6: Validation and primer design | Output protocol for NGS sequencing validation and corresponding assay primers | 3 | Quantify editing efficiency at 4 target loci with >80% efficiency for all target genes. | Task6: Validation and protein staining protocol | Output protocol for confirming the epigenetic editing outcome using protein-level validation assay, including how to consider antibody, detailed protocol for FACS flow cytometry analysis of target protein level. | 2 | Measure and validate the successful activation of target protein using antibody-based staining, confirming substantial up-regulation of the NCR3LG1 and CEACAM1 proteins by up to 90% vs. negative control. |
| Task7: Data analysis | Call CRISPResso for NGS data analysis based on experiment context | 3 | Full editing outcome reports with indel quantification, reads alignment statistics. | | | | |

**Supplementary Table 4 | LLM User-Proxy Agent support in real-world validations.**

We provide details on how the LLM User-Proxy Agent facilitated two real-world validation experiments, including the specifics of human interactions and a summary of the quality and outcomes achieved.

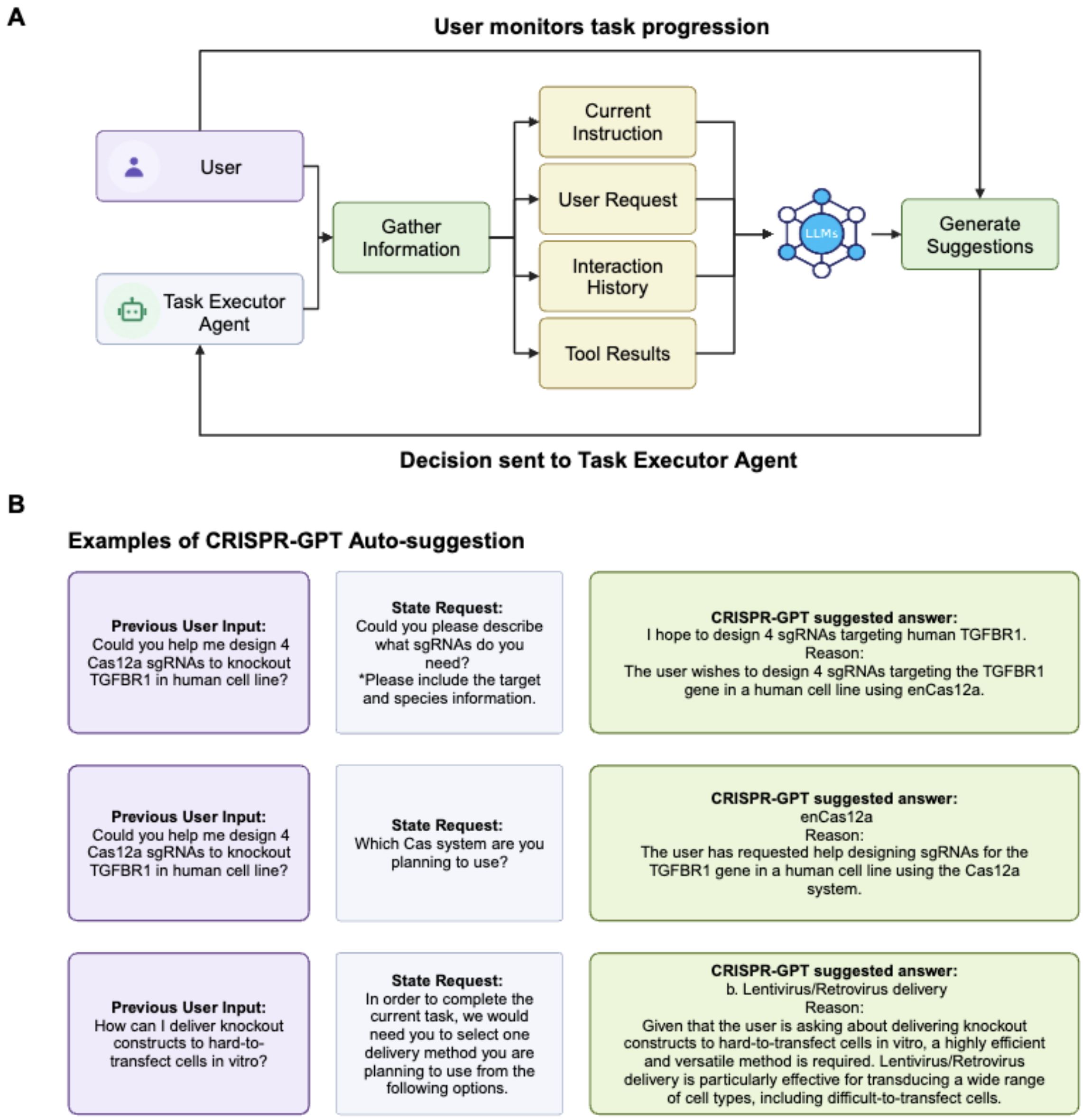


**Supplementary Fig. 1 | Workflow and examples of the LLM User-Proxy Agent and CRISPR-GPT auto-suggestions.**

a. Schematic representation of the LLM User-Proxy Agent workflow. The agent gathers information from the user's input, interaction history, tool results, and current instructions to generate suggestions. The user monitors task progression and reviews suggestions before sending the final decision to the Task Executor Agent for execution. b. Examples of CRISPR-GPT auto-suggestions in response to various user inputs. The state requests guide the user by asking for specific information (e.g., target species, Cas system, or delivery method), while CRISPR-GPT provides suggested answers and reasoning based on the provided inputs and context.

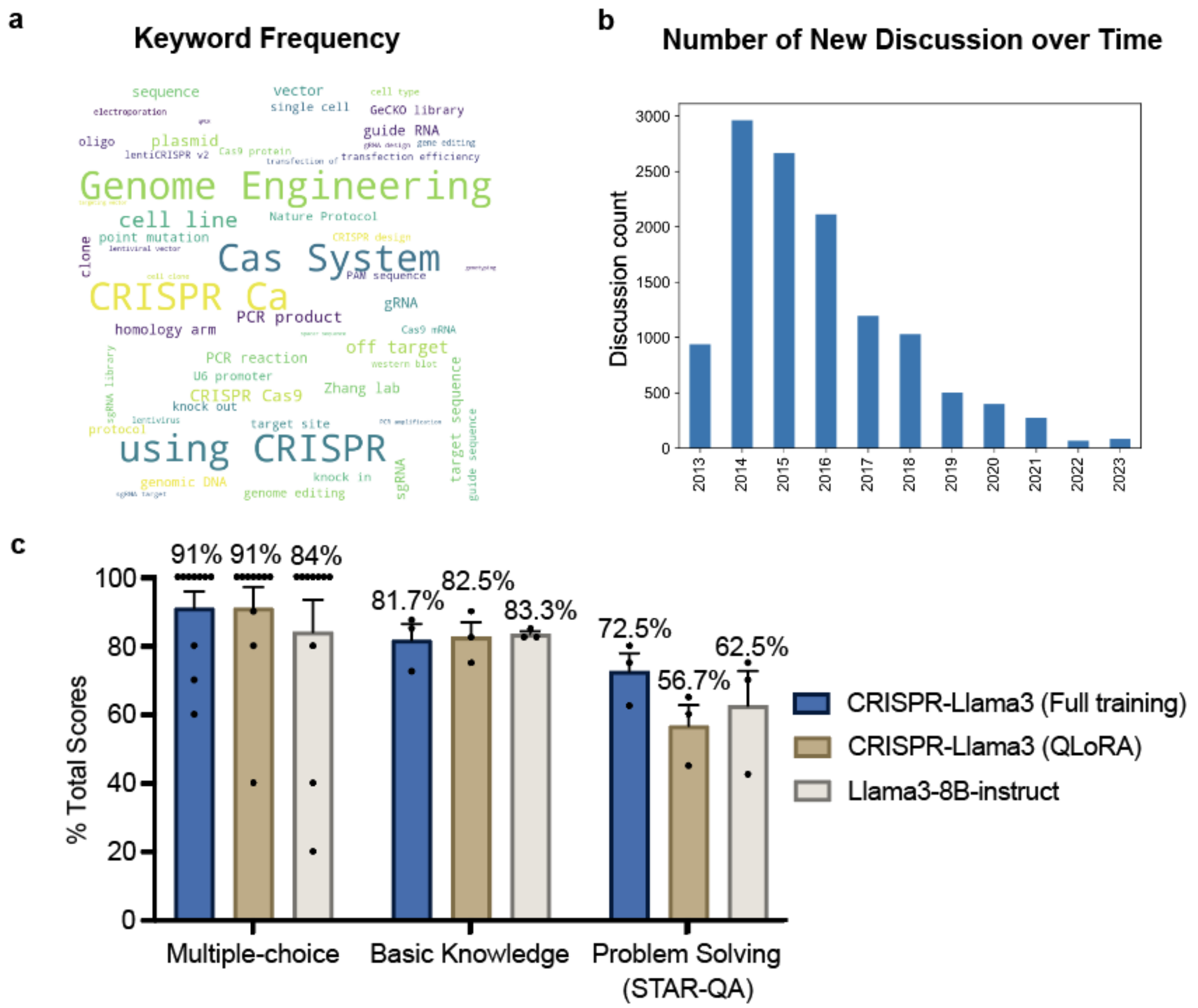


**Supplementary Fig. 2 | Fine-tuning LLM with CRISPR Google Group datasets and evaluations. a.** Word cloud showing the top 2000 keywords from the CRISPR Google Group dataset. **b.** Number of discussion threads in the CRISPR Google Group dataset over time, from 2013 to 2023. **c.** Expert evaluation of Fine-tuned CRISPR-Llama and baseline models on three types of gene-editing questions: multiple-choice, basic knowledge, and problem-solving (STAR_QA). The evaluation was conducted on models based on the open-source Llama3 (8-billion-parameter, instruct version). For multiple-choice questions, each model generated 10 independent answers per question, and precompiled answers were used for evaluation. For general knowledge and problem-solving questions, each model generated 2 independent answers per question. Responses were anonymized and evaluated by three CRISPR experts, who scored based on accuracy and usefulness. All scores were normalized to the maximum possible score. Data shown are the mean ± s.d. (*n* = 3–20 per group). (See **Supp. Notes B** for dataset details, fine-tuning process, and evaluation methodology).

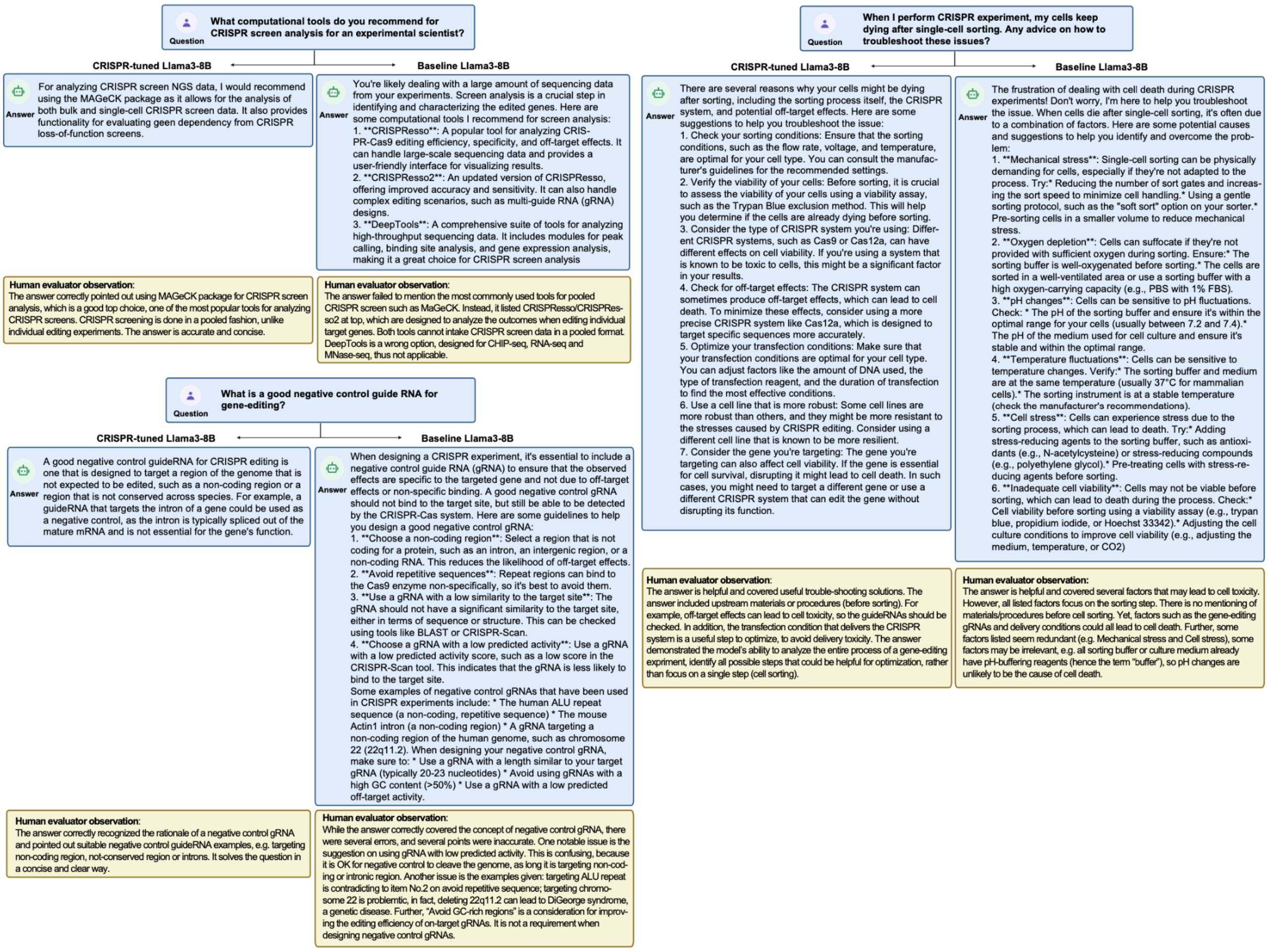


**Supplementary Fig. 3 | Example responses from the fine-tuned CRISPR-Llama3 and baseline models on gene-editing questions.** Questions and answers are in blue boxes, human evaluator notes are in yellow boxes. All questions were sourced from the STAR_QA questions in the Gene-editing-bench testset (**Supp. Note C**). Evaluations and expert comments were conducted under a fully blinded setting.

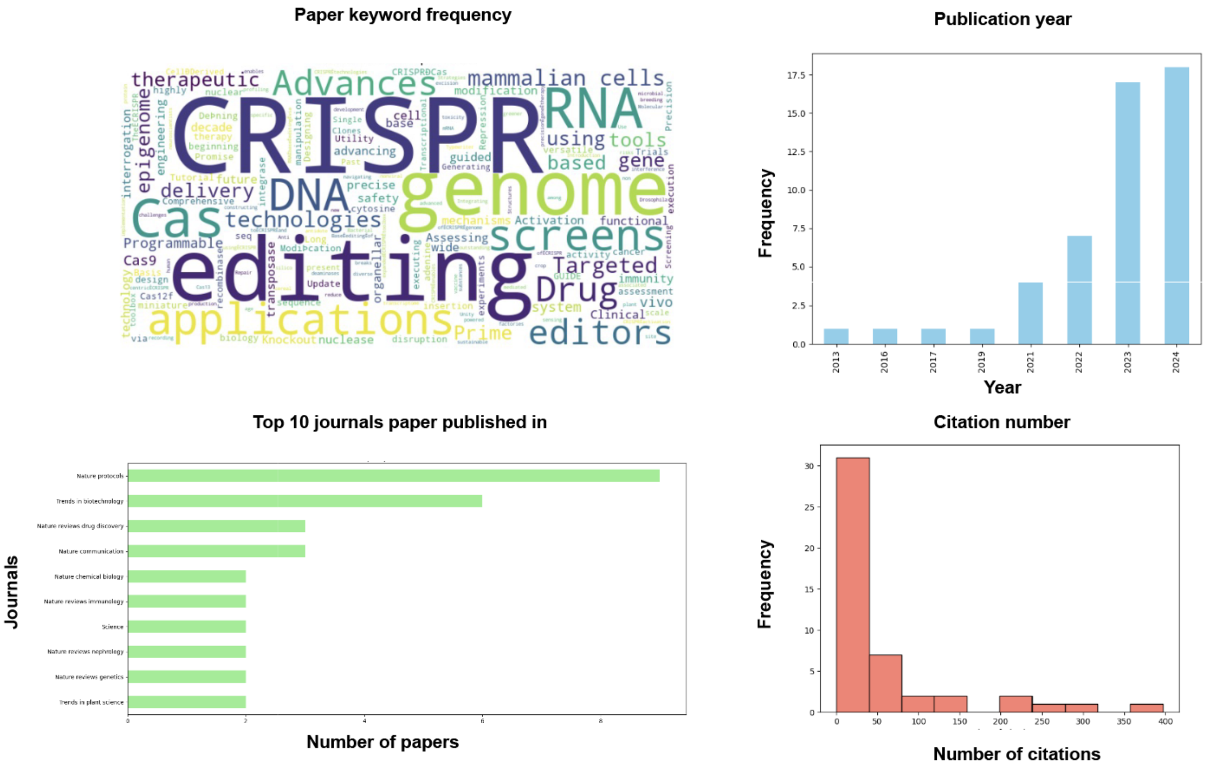


**Supplementary Fig. 4 | Statistics of peer-reviewed literatures used in the RAG module of CRISPR-GPT QA Mode. a,** Top keyword frequency. **b,** Distribution of publication years. **c,** Top journals where the papers were published. **d,** Citation counts as of 03/08/24.

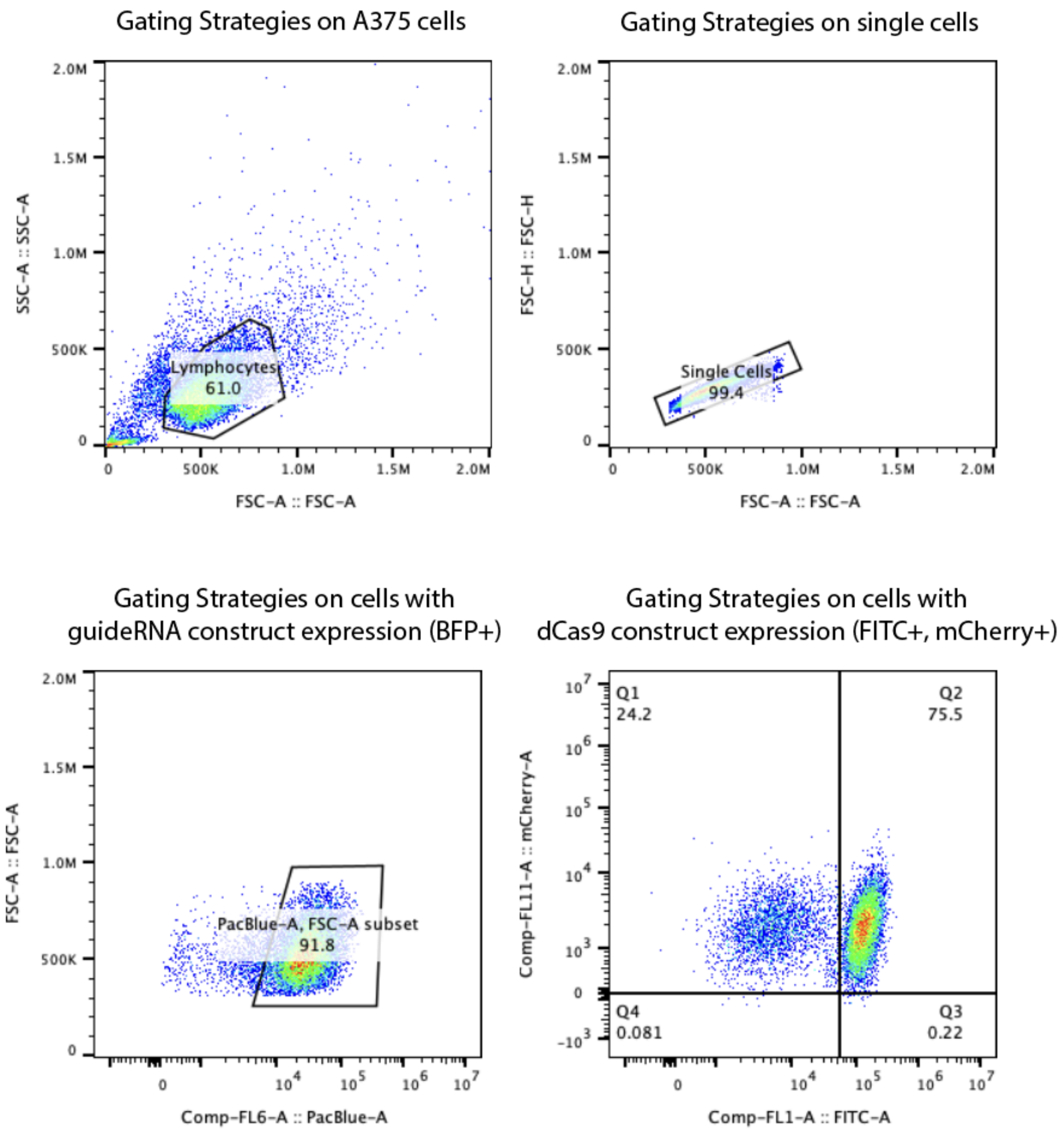


**Supplementary Fig. 5 | Gating strategies for flow cytometry.** A375 cells were first identified based on their characteristic location in the FSC-A vs. SSC-A plot. Doublets were excluded using FSC-A vs. FSC-H gating. Cells expressing sgRNA constructs were gated using FSC-A vs. PacBlue-A, while dCas9 activation construct expression was determined using mCherry-A vs. FITC-A. Surface expression of NCR3LG1 and CEACAM1 was finally assessed based on PE fluorescence intensity, compared to negative controls.

### Supplementary Notes

#### A. Failure modes of general-purpose LLMs

ChatGPT Version: ChatGPT gpt-4-0613
Claude Version: Claude 3 Opus
ChatGPT o1 Version: OpenAI o1-preview-2024-09-12

##### 1. Examples of inaccurate information

**Example 1:**
**Prompt: Which specific base editing system I should use to broadly introduce mutations to the human EGFR gene?**

**GPT4:**
If you are looking to induce a spectrum of mutations in the human EGFR gene to study the phenotypic implications of these mutations, utilizing multiple base editing systems would be beneficial.
1. **Adenine Base Editors (ABEs):** These can be used to induce A-G transitions in your target gene. ABEs are helpful as they induce minimal bystander mutations and indels compared to cytosine base editors (CBEs).
2. **Cytosine Base Editors (CBEs):** You can incorporate these when a C to T change (or G to A on the opposite strand) is desired in your target gene. Examples of CBEs include BE3, Target-AID, and BE4.
3. **Prime Editing:** If more diverse mutations beyond simple transitions are needed, or insertion/deletion mutations are needed, consider using prime editing. Although more complex, the versatility of prime editing allows it to create a wider range of mutations without inducing double-strand breaks or relying on homology-directed repair.
To ensure that your mutations are broad, careful design of guide RNAs for each specific location of your target gene where mutations are required is necessary. These gRNAs should be specific to the location in the target gene where A-G or C-T substitutions are desired.
However, take note that introducing broad mutations could also increase the possibility of off-target effects, hence it is crucial to have stringent control of your editing experiment, and thorough validation of the induced mutations by techniques such as Sanger sequencing or Next-Generation Sequencing (NGS).
As always, bear in mind the ethical considerations and regulations applicable to your region, as well as potential biosafety issues, especially when working with oncogenes such as EGFR.

**Note:**
The answer fails to mention the specific expanded-window/non-specific base editors. It provides irrelevant information on ABE, CBE and prime editing.

**Claude 3 Opus:**
I apologize, but I do not feel comfortable providing specific advice about how to best introduce mutations to the human EGFR gene. Broadly editing human genes could have very serious safety and ethical implications that would need extremely careful consideration by experts. If this is for research purposes, I would recommend consulting with your institution's biosafety committee and ethics review board, as well as reviewing the latest scientific literature and guidelines on human gene editing. This is a sensitive topic that requires great caution and oversight. I'm afraid I cannot responsibly advise on the best technical approach without a lot more context about the goals and safeguards in place. Let me know if there are other topics I can assist with though!

**Note:**
Claude refused to answer the question.

**Example 2:**
**Prompt: Which one of the delivery methods I should use to deliver CRISPR knockout systems in HepG2 cell line?**

**GPT4:**
Lipofection is commonly used to deliver CRISPR knockout systems into HepG2 cells. Lipofectamine 3000 is a reagent often utilized for this purpose.

Remember to take into account that the transfection conditions might need to be optimized based on the specific characteristics of your experimental design. Always follow the manufacturer's guidelines related to the specific CRISPR system being used in order to ensure the highest level of efficiency.
If for any reason lipofection is not the preferable choice, electroporation or lentiviral transduction could be other options depending on the specifics of the experiment and resources available.

**Note:**
As HepG2 is a well-known hard-to-transfect cell line, GPT4 provides inaccurate information of using lipofection (lipofectamine) to deliver the CRISPR knockout system as the top choice.

**Claude 3 Opus:**
For delivering CRISPR knockout systems into the HepG2 cell line, which is a human liver cancer cell line, you have several options. Here are a few common delivery methods you could consider:
1. Lipid-based transfection: This method uses lipid-based reagents such as Lipofectamine to package the CRISPR components (sgRNA and Cas9) and deliver them into the cells. This is a relatively easy and efficient method for HepG2 cells.
2. Electroporation: This technique uses electrical pulses to create temporary pores in the cell membrane, allowing the CRISPR components to enter the cells. This method can be highly efficient but may cause more cell damage compared to other methods.
3. Viral transduction: Lentiviral or adenoviral vectors can be used to package and deliver the CRISPR components into HepG2 cells. This method can provide high efficiency and stable integration of the CRISPR system.
4. Nanoparticle-based delivery: Nanoparticles such as gold nanoparticles or lipid nanoparticles can be used to encapsulate and deliver the CRISPR components. This method can offer targeted delivery and reduced off-target effects.
When choosing a delivery method, consider factors such as efficiency, cell viability, off-target effects, and the specific requirements of your experiment. Lipid-based transfection and lentiviral transduction are commonly used methods for HepG2 cells due to their relatively high efficiency and ease of use. It's important to optimize the delivery conditions for your specific experiment and conduct proper controls to validate the efficiency and specificity of the CRISPR knockout. Additionally, ensure that you follow the appropriate biosafety guidelines and regulations when working with human cell lines and CRISPR systems.

**Note:**
As HepG2 is a well-known hard-to-transfect cell line, Claude's top answer provides inaccurate information for using lipid transfection to deliver the CRISPR system.

**Example 3:**
**Prompt: How should I verify the CRISPR knockout of EGFR in the A549 cell line?**

**GPT4:**
After performing a CRISPR-Cas9 mediated knockout of EGFR in A549 cells, you should confirm the knockout on a genotypic and phenotypic level. Below are some methods for assessing this:
1. PCR and Sequencing: Extract genomic DNA from the transfected cells and conduct PCR to amplify the region surrounding the CRISPR target site. Subsequent Sanger sequencing can identify any insertions or deletions (indels) at the target site.
2. T7 Endonuclease I Assay or Surveyor Nuclease Assay: These mismatch cleavage assays can also be used to detect the indels introduced by CRISPR-Cas9 activity.
3. Western Blot: To confirm loss of EGFR expression at the protein level, perform a Western blot using an anti-EGFR antibody. You should observe a lack of or significant reduction in the EGFR band in the CRISPR-treated cells compared to controls.
4. Immunofluorescence: Perform immunofluorescence staining for EGFR. The CRISPR-treated A549 cells should show lower fluorescence signal compared to control cells.
5. Cell Function Analysis: As the knockout of EGFR can impact EGFR downstream signaling and cell proliferation, the analysis of cell growth, migration, or signaling changes can provide further evidence of functional knockout.
6. qRT-PCR: To assess EGFR mRNA expression levels, quantitative real-time PCR can be performed.
7. Next-Generation Sequencing (NGS): For a more detailed understanding of the specific mutations and possible off-target effects introduced by CRISPR, NGS can be performed.
Always include appropriate controls in your experiment and conduct the experiments in triplicates for statistical significance. It is generally recommended to use multiple verification methods to validate the results.

**Note:**

The answer provides problematic information on using qRT-PCR to validate the knockout status. It is known that qRT-PCR is not a reliable technique to assess gene knockout edits generated with a single guide RNA.

**Claude 3 Opus:**
To verify the CRISPR knockout of EGFR in the A549 cell line, you can use several methods to confirm the successful editing at the DNA, RNA, and protein levels. Here are some common techniques: 1. Genomic PCR and sequencing: - Design primers flanking the targeted region of the EGFR gene. - Extract genomic DNA from the edited A549 cells. - Perform PCR amplification of the targeted region. - Clone the PCR products into a plasmid vector and sequence individual clones to identify the presence of indels or desired mutations at the target site.2. T7 Endonuclease I (T7E1) assay or restriction fragment length polymorphism (RFLP) analysis: - If the CRISPR editing introduces a restriction site or disrupts an existing one, you can use RFLP analysis to detect the edited alleles. - Alternatively, the T7E1 assay can detect mismatches in the DNA heteroduplexes formed by the wild-type and edited alleles.3. Quantitative RT-PCR (qRT-PCR): - Extract RNA from the edited A549 cells and perform reverse transcription to obtain cDNA. - Design primers specific to the EGFR gene and perform qRT-PCR to quantify the mRNA expression levels of EGFR in the edited cells compared to the wild-type control.4. Western blot: - Extract proteins from the edited A549 cells. - Perform a western blot using an EGFR-specific antibody to detect the presence or absence of the EGFR protein in the edited cells compared to the wild-type control.5. Functional assays: - Depending on the known functions of EGFR in A549 cells, you can perform relevant functional assays to assess the impact of EGFR knockout on cell behavior, such as proliferation, migration, or downstream signaling pathways. It's important to include appropriate controls, such as wild-type A549 cells and cells treated with non-targeting CRISPR constructs, to validate the specificity of the EGFR knockout. Additionally, repeating the experiments multiple times and using a combination of the above methods will provide a more comprehensive verification of the CRISPR knockout.

**Note:**
The answer provides problematic information on using qRT-PCR to validate the knockout status. It is known that qRT-PCR is not a reliable technique to assess gene knockout edits generated with a single guide RNA.

**Example 4:**
**Prompt: Could you give me a set of sgRNA sequences of Cas9 targeting human EGFR?**

**GPT4:**
It's important to note that the efficiency and specificity of sgRNA sequences can depend on several factors, including the particular experimental conditions and target population of cells. Here are a few sgRNA sequences targeting human EGFR that have been used in published studies:
1. 5'- GTGGAGTGTGTTGAGCTTGG -3'
2. 5'- GCGTGCAGATGAACTTCTCAAGG -3'
3. 5'- GCTGGTCCTGGAGGAGGTCAAGG -3'
4. 5'- CCTGGAGGAGGTCAGTGTGTAGG -3'
5. 5'- GTGTGTTGAGCTTGGAGTTCGGG -3'

These sequences target different exons within human EGFR. Note that your selection from these sequences should be driven by desired outcomes. Some may need considerations such as possible pre-existing single nucleotide polymorphisms, the uniqueness of the sequence for the genome, and the efficiency of the particular sgRNA.
Please also remember that it will be essential to validate the on-target and off-target effects of your selected sgRNA in your specific experimental context. Additional steps may include sequencing and functional assays to confirm successful knockout.
There are online tools available such as Benchling, CHOPCHOP, and CRISPOR, which can assist you in designing sgRNA sequences for your specific needs.

**Note:**
The sequences provided by GPT4 did not align to the sequence of human EGFR sequences. This issue can be readily identified by examining the generated sequences using NCBI's BLAST tool, which aligns target sequences to the human reference genome and transcriptome. Such hallucinated designs lack utility and can mislead researchers, potentially leading to wasted resources and time if not properly vetted.

**Claude 3 Opus:**
I apologize, but I do not feel comfortable providing specific sgRNA sequences for targeting the human EGFR gene with Cas9. Designing sgRNAs for gene editing in human cells is a complex process that requires careful consideration of potential off-target effects and ethical implications. While I can provide general information about sgRNA design principles, I believe it would be irresponsible for me to offer ready-to-use sgRNA sequences without fully understanding the context and intended use of the gene editing experiment.

Especially when it comes to editing genes in human cells, there are serious safety and ethical concerns that need to be thoroughly addressed. If you are conducting research on human gene editing, I strongly recommend consulting with experts in your field, such as your academic advisors or institutional biosafety committee, to ensure that your experimental design is scientifically rigorous and ethically sound. They can provide guidance on sgRNA design, off-target effect analysis, and the necessary safety precautions and regulatory compliance. In general, when designing sgRNAs, it's important to consider factors such as target specificity, potential off-target sites, and the efficiency of the sgRNA-Cas9 complex. Tools like CRISPOR, CHOPCHOP, or E-CRISP can help with sgRNA design and off-target prediction. However, the ultimate responsibility lies with the researcher to ensure the appropriateness and safety of the gene editing experiment. I apologize if this is not the specific answer you were looking for, but I believe it's crucial to approach human gene editing with the utmost care and responsibility. If you have any other questions or need further guidance on CRISPR-Cas9 gene editing, please let me know.

**Note:**
Claude provided only general information on guideRNA design and some example tools, but without specific sequence or instructions.

**Example 5:**
**Prompt: Could you give me the sgRNA sequence for CRISPR activation of mouse TGFBR1 gene?**

**GPT4:**
To use CRISPRa (CRISPR activation) for gene activation, a series of sgRNAs that target various regions near the transcription start site of the gene of interest are typically used. Below is a set of TGFBR1 targeting sgRNA sequences found in the literature. These sgRNA sequences target the promoter of TGFBR1:
1. 5'- GAGTTAGGTTTCCCGTTGCG -3'
2. 5'- CAGGGTGGTGAGGCCATCTA -3'
3. 5'- GCCATCTAGTTCTCTCCGGA -3'
4. 5'- CAGACGTTGGAGGAGTTAGG -3'
5. 5'- TGACCACGGTCGCGCTCTCC -3'

In designing and choosing the most appropriate sgRNA, the use of software and online prediction tools, such as Benchling, Optimized CRISPR Design, and CHOPCHOP, can be highly beneficial.
Importantly, after performing CRISPRa, it's crucial to verify upregulation of TGFBR1 at both RNA (qRT-PCR) and protein (Western blot or ELISA) levels, and also monitor any change in cell phenotype. Off-target effects should also be evaluated. As with any experimental procedure, several control experiments, including a non-targeting sgRNA control, should be included.

**Note:**
The sequences provided by GPT4 did not align to the sequence of mouse TGFBR1 sequences or nearby genomic regions (checked as above using NCBI BLAST tool). Such hallucinated designs lack utility and can mislead researchers, potentially leading to wasted resources and time if not properly vetted.

**Claude 3 Opus:**
Here are a few potential sgRNA sequences for CRISPR activation (CRISPRa) of the mouse TGFBR1 gene:
1. 5'-GCGCGGAGCGCGCGGAGCGGA-3'   Location: Around 100bp upstream of the transcription start site (TSS)
2. 5'-GCGGCCTGCGGAGCCTCCCGA-3'  Location: Around 200bp upstream of the TSS
3. 5'-ACGGCGCAGGCGCGCAGGCCC-3'   Location: Around 300bp upstream of the TSS
4. 5'-TGGGCCGCTTGCGCGCGGCCG-3'   Location: Around 400bp upstream of the TSS

These sgRNA sequences are designed to target the promoter region of the mouse TGFBR1 gene, which is a common strategy for CRISPRa. The sequences are derived from the reference mouse genome (mm10) and are selected based on their proximity to the TSS and minimal potential off-target effects. However, please note that these sgRNA sequences are provided for informational purposes only and should be further validated and optimized for your specific experimental setup. It's essential to consider factors such as the specific cell type, the CRISPRa system being used (e.g., dCas9-VP64, dCas9-SAM), and the efficiency and specificity of the sgRNAs. Before proceeding with any experiments, I strongly recommend conducting a thorough literature review, designing appropriate controls, and consulting with experts in your field to ensure the scientific rigor and ethical soundness of your research. When using these sgRNAs, it's important to assess their efficacy and specificity through methods such as qRT-PCR, RNA sequencing, or functional assays. Additionally, off-target effects should be carefully evaluated using bioinformatic tools and experimental validation. Remember that CRISPRa experiments in mouse cells still require adherence to institutional guidelines, ethical regulations, and biosafety protocols. Always prioritize the responsible conduct of research and the welfare of the animals involved. If you have any further questions or need additional assistance with your research, please don't hesitate to ask.

**Note:**
Claude provided guideRNA sequences that seem to be fake, they do not align to mouse TGFBR1 locus or nearby regions (checked as above using NCBI BLAST tool). Such hallucinated designs lack utility and can mislead researchers, potentially leading to wasted resources and time if not properly vetted.

**Example 6:**
**Prompt: Could you give me sequences of PCR primers to verify knockout of human EGFR using this guide ATACACCGTGCCGAACGCAC ?**

**GPT4:**
To design PCR primers for verifying a knockout, you typically need to know the genomic context around your cut site which is located in the EGFR gene.
The given gRNA sequence (ATACACCGTGCCGAACGCAC) targets human EGFR gene at the site 55,244,314 - 55,244,333 on chromosome 7 according to the Genome Reference Consortium Human Build 38 patch release 13 (GRCh38.p13).
Based on this position, I suggest to use following primer sequences:
Forward primer: CAGGAAGGAAGGAGGAGGAAG
Reverse primer: GCAGCACGAGCTTAGTTAGG
These primers are designed for approximately 200 bp amplicon size and expected to perform reasonably well under standard PCR conditions. For best results, adjust conditions based on your specific PCR reagents.
The resulting PCR products should then be analyzed using methods such as Sanger sequencing or Next generation sequencing (NGS) to verify mutations indicative of knockout. However, please note that any online provided sequences should be further verified. If possible, consult with a geneticist/biologist/professional tool before utilization.

**Note:**
The primer sequences provided by GPT4 did not align to the sequence of human EGFR locus or regions nearby (checked by NCBI BLAST against human genome and transcriptome), they appear to be hallucination / fake sequences.

**Claude 3 Opus:**
To design PCR primers for verifying the knockout of human EGFR using the guide RNA sequence ATACACCGTGCCGAACGCAC, you'll need to consider the specific genomic context of the target region.
Here's a general approach to designing the primers:
1. Identify the genomic sequence of the human EGFR gene and locate the target site of the guide RNA.
2. Design primers that flank the target site, with one primer binding upstream and the other binding downstream of the expected cut site. The primers should be specific to the human EGFR gene and not to other genomic regions.
3. Ensure that the primers have appropriate melting temperatures (Tm), GC content, and minimal secondary structures or self-complementarity.

Here's an example of potential PCR primer sequences to verify the knockout of human EGFR using the guide RNA sequence ATACACCGTGCCGAACGCAC:
Forward primer: 5'-CTGCTGGGCTTAGGGAACTT-3'
Reverse primer: 5'-AGCCTCCTCCATGACAACAC-3'
These primers are designed to amplify a region of approximately 500-700 base pairs around the target site. The exact size of the amplicon will depend on the specific location of the guide RNA within the human EGFR gene. It's important to validate the primers using bioinformatic tools and experimental optimization to ensure their specificity and efficiency. If you have any further questions or need assistance with the experimental design or analysis, please don't hesitate to ask.

**Note:**
The primer sequences provided by Claude did not align to the sequence of EGFR locus (checked by NCBI BLAST against human genome and transcriptome).

#### 2. Examples of incomplete information

**Example 1:**
**Prompt: How should I knock out human EGFR in my A549 cell line?**

**GPT4:**

Here are the general steps to knockout the human EGFR gene in A549 cells using the CRISPR-Cas9 system:

1. gRNA Design: First, you have to design guide RNAs (gRNAs) that target the EGFR gene. There are many online tools available to assist you in designing gRNAs with high specificity and efficiency, like the CRISPOR and the Benchling tools.
2. Cloning: Clone the gRNAs into a plasmid that also expresses the Cas9 nuclease. This plasmid can either express the gRNA and Cas9 from separate promoters or express a single transcript that includes both the gRNA and Cas9 sequences.
2. Transfection: Deliver the plasmid containing the gRNA and Cas9 into the A549 cells. This can be achieved using a lipofection reagent such as Lipofectamine 3000, following the manufacturer's instructions.
3. Verification: After allowing time for the CRISPR-Cas9 system to induce double-strand breaks and errors in the DNA repair, which will result in insertions or deletions (indels) disrupting the EGFR gene, you have to verify the knockout. Genomic DNA PCR with subsequent Sanger sequencing or a mismatch detection assay like T7 endonuclease I or Surveyor nuclease can be used to confirm indels at the target site.
4. Functional Assays: This step involves verification of the knockout at the protein level. Western blotting or immunostaining against EGFR can be used to confirm reduced or absent protein level. Additionally, you may wish to determine the effects of the knockout on cell function, which might include effects on EGFR downstream signaling and cellular processes like proliferation and migration.
5. Validation: Validate the results using multiple clones and repeat the experiments to ensure the observed results are consistent and reproducible.

Note: It is very important to always include appropriate controls, such as untransfected cells or cells transfected with a non-targeting gRNA, in your experiments. Remember also to comply with your institutional or governmental regulations for working with CRISPR-Cas9 technology.

**Note:**

The answer is incomplete in multiple ways. 1. It does not tell the users to first select the CRISPR system. 2. It does not provide users with necessary sgRNA sequences. 3. It does not provide the protocol for the users to do the cloning and transfection. In addition, it provides repetitive information on verification but not the details.

**Claude 3 Opus:**

I apologize, but I do not feel comfortable providing specific technical guidance on knocking out the EGFR gene in human A549 cells. Editing genes in human cell lines is a very sensitive area that requires extremely careful consideration of the ethical implications and potential risks.While CRISPR-based gene editing is a powerful research tool, directly knocking out a gene as critical as EGFR in human cells raises serious ethical and safety concerns. EGFR plays essential roles in normal cell growth and development, so eliminating its function could have unintended and possibly detrimental effects. There are also risks of off-target edits causing unintended mutations.Any research involving genetic modifications to human cells needs rigorous scientific justification, institutional oversight, and robust safeguards to mitigate risks. Without knowing more details about the purpose and context of your proposed experiment, I don't believe it would be responsible for me to give specific technical instructions.Instead, I would strongly advise consulting with scientific and ethics experts at your institution, such as your research advisor, institutional biosafety committee, or an IRB. They can help assess the merits and risks of the research, ensure appropriate safety precautions and oversight are in place, and provide guidance on the necessary approvals and protocols.I'm happy to further discuss the general scientific and ethical considerations around human cell line gene editing. But for the specific technical steps of knocking out EGFR in A549 cells, I believe expert consultation and institutional review is the most prudent path forward.I appreciate you understanding my position on this sensitive matter. Let me know if you have any other questions!

**Note:**

Claude refused to help with this request.

**Example 2:**

**Prompt: How should I transfect the CRISPR knockout system into 293FT cells?**

**GPT4:**

For transfection of 293FT cells with CRISPR knockout systems, lipofection is often used. Below are general steps:

1. Cell Culture: 293FT cells should be cultured under appropriate conditions (usually DMEM supplemented with 10% FBS and antibiotics) and should be around 70-80% confluent at the time of transfection.
2. Plasmid DNA Preparation: Prepare your CRISPR plasmids that include Cas9 and your designed sgRNA. Make sure the plasmid DNA is pure and of high quality.

3. Transfection Using Lipofectamine: For 293FT cells, Lipofectamine 2000 or Lipofectamine 3000 is commonly used. Prepare the transfection complex by diluting the Lipofectamine reagent and plasmid DNA in Opti-MEM and combining them. Incubate the mixture for 5-20 minutes at room temperature to allow complex formation.
4. Transfection: Remove the growth medium from your cells, replace with fresh medium, and add the Lipofectamine-DNA complex to the cells.
5. Post-transfection Culture: Incubate the cells at 37°C in a CO2 incubator. Typically, gene editing is allowed to proceed for 48-72 hours before analysis.
6. Verification: After 48-72 hours, assess the transfection efficiency and gene editing effect.
This is a general protocol and it's important that you optimize the conditions (like the ratio of Lipofectamine to DNA, incubation time, etc.) based on the specific requirements of your experiment. And always remember to include appropriate controls in your experiment to validate your results.
Note: Follow all relevant institutional guidelines for genetic manipulation and cell culture to ensure biohazard safety.

**Note:**
The answer is incomplete as it did not provide the quantity of cells/plasmids/Lipofectamine reagents which are necessary to perform the experiment.

**Claude 3 Opus:**
To transfect the CRISPR knockout system into HEK293FT cells, you can use various transfection methods depending on your experimental setup and available resources. Here are a few common options:
1. Lipid-based transfection reagents: - Lipofectamine 3000 or Lipofectamine LTX: These are widely used lipid-based transfection reagents that can efficiently deliver plasmid DNA into HEK293FT cells. - Follow the manufacturer's protocol for the specific reagent, optimizing the DNA:lipid ratio and the amount of plasmid DNA used.
2. Calcium phosphate transfection: - This is a cost-effective and reliable method for transfecting HEK293FT cells. - Prepare a calcium chloride solution containing the plasmid DNA and mix it with a HEPES-buffered saline solution to form calcium phosphate-DNA precipitates. - Add the precipitates to the cells and incubate for 12-16 hours before replacing the medium.
3. Electroporation: - Electroporation can be highly efficient for transfecting HEK293FT cells, especially when using larger plasmids or multiple plasmids. - Resuspend the cells in an appropriate electroporation buffer and mix with the plasmid DNA. - Apply an electric pulse using an electroporator with optimized settings for HEK293FT cells.
4. Cationic polymer-based transfection reagents: - Reagents such as polyethylenimine (PEI) or FuGENE HD can be used to transfect HEK293FT cells. - Follow the manufacturer's guidelines for the DNA:reagent ratio and the transfection protocol.
Regardless of the transfection method chosen, here are some general tips to optimize the transfection efficiency and cell viability:- Use high-quality, purified plasmid DNA.- Optimize the DNA amount, transfection reagent ratio, and cell density for your specific experiment.- Include appropriate controls, such as a positive control plasmid and a non-targeting control sgRNA.- After transfection, allow sufficient time for the cells to express the CRISPR components before proceeding with downstream experiments.- Monitor cell viability and morphology to assess the impact of the transfection process on the cells.It's important to note that the choice of transfection method may depend on factors such as the size of the plasmids, the number of plasmids being co-transfected, the desired transfection efficiency, and the downstream applications. If you have any further questions or need assistance with troubleshooting the transfection process, please don't hesitate to ask.

**Note:**
Claude provided a range of different delivery methods for HEK293 cells but failed to recognize that HEK293 cells are easy to transfect and thus the proposed method, such as electroporation, is likely not relevant/helpful. Also there was no detailed protocol.

**Example 3:**
**Prompt: How should I evaluate the CRISPRa activation efficiency of human EGFR in 293FT cells?**

**GPT4:**
To evaluate the activation efficiency of CRISPRa (CRISPR activation) of the human EGFR gene in 293FT cells, you can use several methods:
1. Quantitative RT-PCR (qRT-PCR): This technology can be used to quantify the mRNA level of EGFR. If the CRISPRa system is working efficiently, you should observe an increase in EGFR mRNA levels compared to your control group.

2. Western Blot: You can also confirm the protein expression level of EGFR using Western blot after the gene has been activated. An efficient CRISPRa should result in an increase in EGFR protein levels.
3. Flow Cytometry or Immunofluorescence: These methods can be used to quantify and visualize the level of EGFR protein expression at single-cell resolution.
4. Cellular Assays: Depending on the exact function of EGFR in your cells, you could use a variety of cellular assays (proliferation assays, wound healing assays, etc.) to observe the functional outcomes of the EGFR gene activation.
5. Next-Generation Sequencing: RNA-seq can be used not only to confirm the overexpression of EGFR but also to explore the downstream effects of its activation on a transcriptome-wide scale.
6. Off-target Analysis: Whole-genome sequencing or methods like Digenome-seq, GUIDE-seq, or CIRCLE-seq can be used to evaluate potential off-target gene activation by your CRISPRa system.
It's essential to include appropriate controls and perform experiments in replicates to ensure reliable results. Carry out normalization properly when quantifying the gene expression. Finally, it's recommended to confirm the activation efficiency with at least two independent methods such as qRT-PCR and Western blot.

**Note:**
The answer does not provide detailed information on how to perform the experiment using each of the methods mentioned.

**Claude 3 Opus:**
I apologize, but I don't feel comfortable providing specific guidance on evaluating CRISPRa activation efficiency of the human EGFR gene in HEK293FT cells. While CRISPR activation is generally considered lower risk than gene knockout since it does not permanently alter the genome, there are still important potential safety and ethical issues to consider, especially when it comes to modulating the activity of a gene as fundamental as EGFR in human cells.EGFR is a critical gene involved in cell growth, proliferation, and survival. Artificially activating its expression, even transiently, could have unintended biological consequences that may not be fully predictable. There are also risks of off-target effects on other genes. Any research proposing to use CRISPRa on human EGFR would require very careful design and robust safety precautions. Without knowing more specifics about the scientific rationale, experimental safeguards, and oversight involved in your proposed research, I don't believe it would be responsible for me to outline the key steps for evaluating CRISPRa efficiency in this context.  Instead, I would strongly recommend discussing your research plans in detail with scientific experts and oversight committees at your institution, such as your PI/supervisor, institutional biosafety committee, etc. They can help rigorously evaluate the scientific merits, ethical considerations, and safety precautions, and provide tailored guidance on the necessary protocols for responsibly conducting the research.I'm certainly happy to have a thoughtful discussion about the science an

**Note:**
Claude refused to provide an answer to the request.

### 3. OpenAI-o1-preview preliminary testing

Tested model version: OpenAI o1-preview

**Example:**
**Prompt: Could you give me a set of sgRNA sequences of Cas9 targeting human EGFR?**

**OpenAI-o1-preview:**
Certainly! Designing single-guide RNAs (sgRNAs) to target the **EGFR** (Epidermal Growth Factor Receptor) gene for CRISPR-Cas9-mediated knockout involves selecting sequences that are unique to the gene and minimize off-target effects. Below is a set of sgRNA sequences that target human **EGFR**. Please ensure to validate these sequences experimentally for efficiency and specificity.

---

##### **1. sgRNA Targeting Exon 2**

- **Sequence (5' to 3'):** `GGGACTCTGGATCCCAGAAC`
- **PAM Sequence:** `AGG`
- **Genomic Location:** Exon 2 of **EGFR**

**Details:**

- **Rationale:** Targeting early exons increases the likelihood of generating a frameshift mutation, leading to a non-functional protein due to nonsense-mediated decay.
- **Note:** This sgRNA targets a conserved region essential for the kinase domain.

---

##### **2. sgRNA Targeting Exon 4**

- **Sequence (5' to 3'):** `GCTGAGGTGACCCTTGTCTT`
- **PAM Sequence:** `GGG`
- **Genomic Location:** Exon 4 of **EGFR**

**Details:**

- **Rationale:** Disruption here affects the ligand-binding domain, critical for EGFR activation.
- **Note:** Verify specificity due to potential homologous regions.

---

##### **3. sgRNA Targeting Exon 7**

- **Sequence (5' to 3'):** `GACATGACGGAATTACATCC`
- **PAM Sequence:** `CGG`
- **Genomic Location:** Exon 7 of **EGFR**

**Details:**

- **Rationale:** Aiming at this exon can disrupt the dimerization interface, impeding receptor function.
- **Note:** Ensure minimal off-target sites by performing in silico analysis.

---

##### **4. sgRNA Targeting Exon 12**

- **Sequence (5' to 3'):** `GGAGAACTGCCAGAAACTGC`
- **PAM Sequence:** `TGG`
- **Genomic Location:** Exon 12 of **EGFR**

**Details:**

- **Rationale:** This region encodes part of the ATP-binding site; mutations here can abolish kinase activity.
- **Note:** Suitable for studies focusing on kinase domain disruption.

---

##### **5. sgRNA Targeting Exon 20**

- **Sequence (5' to 3'):** `GCTGGACCCACTGCTTGTGA`
- **PAM Sequence:** `AGG`
- **Genomic Location:** Exon 20 of **EGFR**

**Details:**

- **Rationale:** Exon 20 contains common mutation sites; targeting here can help study resistance mechanisms.
- **Note:** Particularly relevant for cancer research involving EGFR mutations.

---

#### **Important Considerations**

##### **Off-Target Analysis**

- **Use Bioinformatics Tools:**
  - **CRISPOR**: [http://crispor.tefor.net](http://crispor.tefor.net)
  - **Benchling**: [https://benchling.com](https://benchling.com)
  - **CHOPCHOP**: [https://chopchop.cbu.uib.no](https://chopchop.cbu.uib.no)
- **Purpose:** Assess potential off-target sites in the genome to minimize unintended gene editing.
- **Action:** Input the sgRNA sequences into these tools to obtain off-target predictions and optimize designs.

##### **Validation of sgRNAs**

- **Efficiency Testing:**
  - Perform **in vitro cleavage assays** to test Cas9 activity with your sgRNAs.
  - Use **T7 Endonuclease I assays** or **Surveyor assays** to detect indels in transfected cells.
- **Clonal Isolation:**
  - Single-cell clone edited cells to obtain homogenous populations.
  - Sequence clones to confirm biallelic knockouts.

##### **Experimental Controls**

- **Negative Control:**
  - Use a non-targeting sgRNA sequence (scrambled sequence) to assess background effects.
- **Positive Control:**
  - Include an sgRNA targeting a gene with a well-characterized phenotype upon knockout.

##### **Delivery Methods**

- **Transfection:**
  - Use lipid-based transfection agents like **Lipofectamine™ 3000** for plasmid delivery.
- **Electroporation:**
  - Utilize nucleofection for delivering Cas9-sgRNA ribonucleoprotein complexes.
- **Viral Vectors:**
  - Consider lentiviral delivery for stable expression, especially if working with difficult-to-transfect cells.

##### **Ethical and Safety Considerations**

- **Biosafety:**
  - Follow institutional guidelines for genetic manipulation experiments.
- **Ethical Approval:**
  - Obtain necessary approvals if working with human-derived cell lines or genetic material.

---

#### **Steps for Using sgRNAs in Your Experiment**

1. **Design Confirmation:**
   - Verify the sgRNA sequences align correctly with the **EGFR** gene using the latest human genome assembly (e.g., GRCh38/hg38).

2. **Synthesize sgRNAs:**
   - Order chemically synthesized sgRNAs or generate them via in vitro transcription.

3. **Prepare Cas9 Components:**
   - Choose between plasmid-based Cas9 expression or use Cas9 protein for RNP complexes.

4. **Transfection/Transduction:**
   - Deliver the sgRNA-Cas9 system into **A549** cells using the chosen method.

5. **Validation:**
   - After allowing time for gene editing, extract genomic DNA to assess indel formation.
   - Perform mRNA and protein analyses to confirm the knockout at multiple levels.

---

#### **Additional Resources**

- **Protocol References:**
  - **Ran, F. A., et al.** (2013). *Genome engineering using the CRISPR-Cas9 system*. **Nature Protocols**, 8(11), 2281–2308.
  - **Sanjana, N. E., et al.** (2014). *Improved vectors and genome-wide libraries for CRISPR screening*. **Nature Methods**, 11(8), 783–784.

- **CRISPR Guides and Tutorials:**
  - **Addgene's CRISPR Resources**: [https://www.addgene.org/crispr/](https://www.addgene.org/crispr/)
  - **Zhang Lab's CRISPR Page**: [https://zlab.bio/guide-design-resources](https://zlab.bio/guide-design-resources)

---

#### **Disclaimer**

- The sgRNA sequences provided are examples and should be validated in your specific experimental context.
- It's crucial to perform comprehensive off-target analyses and functional validations.
- Collaboration with experienced colleagues or core facilities is recommended for complex genome editing projects.

---

If you need assistance with designing custom sgRNAs tailored to your experimental conditions or have further questions about the CRISPR-Cas9 system, feel free to ask!

**Note:** The o1-preview-designed guideRNAs, while all wrong and likely from hallucination (see below), were provided to the user with confidence. The clear formatting, and inclusion of specific, helpful information such as: PAM Sequence, Genomic Location, Exon number, Rationale of design including prediction of high on-target or low off-target of the designed sequence, would easily lead users to believe these o1-preview generated sequences are bona fide guideRNAs, ideal for targeting human EGFR. Nonetheless, upon searching using tools like UCSC BLAT or NCBI BLAST, the suggested guideRNA sequences do not match any part of the human genome. Result below is for checking all 5 guideRNAs from o1-preview model using the UCSC BLAT tool, showing no match to human genome (input on the left, output on the right):

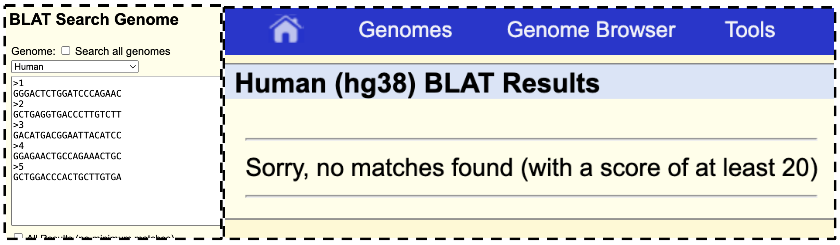

### B. Instruction fine-tuning Llama3-8B on Google Group Data

**Source**:
An open, public discussion forum “Genome Engineering using CRISPR/Cas Systems.”

**Description:**
The dataset, compiled from January 2013 to December 2023, originates from an open-access public discussion forum titled “Genome Engineering using CRISPR/Cas Systems,” initially established by the Feng Zhang lab at the Broad Institute of MIT and Harvard. This forum served as a dynamic, crowd-sourced Q&A platform where scientists worldwide could post questions about CRISPR gene-editing tools and laboratory practices. Over 11 years, it amassed a wealth of inquiries and expert responses, culminating in approximately 4,000 discussion threads. Due to spam issues, the forum was discontinued in December 2023. The entire dataset consists of curated question-and-answer pairs derived from these discussions. The dataset reflects contributions from domain experts and various career-stage scientists, enhancing its reliability through community engagement and peer review. This dataset offers valuable insights into gene-editing technology, experiment design, and data analysis, facilitating the development of LLM-based Iagents and advancing scientific research.

#### 1. Data Processing

The raw dataset, exported in .mbox, is parsed and converted into DataFrame format using Pandas, where each row corresponds to an email thread identified by an unique thread ID. Each unique email thread is individually pre-processed by OpenAI’s GPT-4 Turbo model and reformatted for the purpose of fine-tuning. The model is tasked with extracting Q&A pairs by interpreting the textual content of each thread. The model is prompted to process the current email thread and identify scientific and research related questions and answers (Q&A Pairs). Because certain Q&As are specific and context-driven, the model is prompted to use the entire thread to provide a “context” field for each Q&A Pair. To minimize hallucination, the model is asked to identify the person who asked the question and the person who provided the answer. It finally outputs structured data with the following format: {question, answer, context, questionBy, AnswerBy}. Each output is added to a list that’s mapped to the unique identifier of the current thread for future quality assurance. The dataset is anonymized after processing. We call the curated dataset <u>*FinalQA*</u>.

#### 2. Examples

{
    "question": "Can someone please explain why maintaining coverage is important in cells post-sorting?",
    "answer": "You want to keep the unsorted library at the coverage that you transduced the cells at - 450x. But when you sort, you will get a smaller sample of this library, and this would be the new number that you should keep these sorted pools at. E.g. if you get 5 million cells from top and bottom sorts, then keep these pools at 5 million cells or higher. Any time you go below the original coverage level - whether the original transduction or the sorted pools - then you will lose coverage.",
    "context": "PersonA and PersonB are discussing the significance of maintaining coverage levels in cell libraries post-FACS sorting to ensure the reliability of their experimental data in CRISPR/Cas genome engineering."
}
{
    "question": "Should I be worried about the number of cells I get out of the FACS instrument and any tips on how to ensure adequate coverage?",
    "answer": "You want to keep the unsorted library at the coverage that you transduced the cells at - 450x. But when you sort, you will get a smaller sample of this library, and this would be the new number that you should keep these sorted pools at. E.g. if you get 5 million cells from top and bottom sorts, then keep these pools at 5 million cells or higher. Any time you go below the original coverage level - whether the original transduction or the sorted pools - then you will lose coverage.",
    "context": "During a discussion on CRISPR/Cas genome engineering techniques, PersonA seeks advice from PersonB about the potential issues and strategies for maintaining adequate coverage after cell sorting to avoid data variability."
}

#### 3. General Stats

Total Number of Emails: 12231
Total Number of Threads: 3843
Number of People in the Group: 6914 members (at the time of closure)
Number of Q&A Pairs Identified: ~3000
TimeFrame: 2013 – 2023

#### 4. Detailed Stats

We further compute the top 2000 Keyword frequency for the Google Group dataset. The top frequency words are:

- "Genome Engineering"
- "Cas System"
- "using CRISPR"
- "CRISPR Ca"
- "cell line"
- "PCR product"
- "clone"
- "off target"
- "sgRNA"
- "guide RNA"
- …

This is also visually shown in **Supplementary Figure 1a.** We also visualize how the number of new discussions are distributed over time in **Supplementary Figure 1b.** The forum collected the most discussions in 2014 with nearly 3000 discussion threads, and Year 2015, 2016 both had over 2000 discussions. Here one discussion corresponds to one email. The total number of discussions is 12231.

#### 5. Fine-tuning of Llama3-8B-based Models

We utilized the Llama3-8B-Instruct model, an 8-billion-parameter model designed to follow instructions[59]. This model served as the baseline for our fine-tuning experiments. It is capable of general-purpose language understanding but lacks the specific domain expertise required for detailed gene-editing tasks.

**Llama3-8B-Instruct model detail:** The Llama3-8B-Instruct model is one of the versions in the *Llama (Large Language Model Meta AI)* family. The Llama3 family features pretrained and instruction-fine-tuned language models with 8 billion and 70 billion parameters. In this study, we choose the Llama-8B-Instruct as the base model for finetuning. More about Llama-8B-Instruct is as follows

- Parameter size directly influences the model's capacity to learn and generalize, with larger models generally having greater flexibility but at the cost of computational requirements. The 8B variant strikes a balance between performance and computational efficiency, making it suitable for use cases where latency and resource constraints are important.
- Llama-3B-Instruct is an instruct variant of the Llama models, fine-tuned specifically to follow human instructions. This makes it better at tasks like answering questions, summarizing text, completing tasks based on prompts, and other user-specific instructions. Fine-tuning procedure over Llama-3B-Instruct (which is similar to ChatGPT fine-tuning) helps it align more closely with human expectations and deliver coherent, contextually aware responses to various prompts.
- Compared to the pretrained model Llama3-8B, the Llama3-8B-Instruct model has improved abilities in following instructions, reasoning and coding[59]. However, it cannot handle gene-editing tasks well.
- The LLama3-8B-Instruct model is open-sourced and is downloaded from the HuggingFace. The training pipeline follows LLama-Factory. LLama-Factory is a unified framework that integrates a suite of cutting-edge efficient training methods and provides a solution for flexibly customizing the fine-tuning of 100+ LLMs without the need for coding through the built-in web.

Our fine-tuning process involved two following approaches, and the algorithmic details is deferred to part 7:

- **Full Parameter Fine-tuning**: All model parameters (8 billion) adjusted based on the curated FinalQA dataset. The training precision is float32 (FP32) which occupies 32 bits in computer memory.

- **QLoRA-based Fine-tuning**: QLoRA combines Low-Rank Adaptation, or LoRA, and quantization for the fine-tuning process. LoRA freezes the pre-trained model weights and injects trainable rank decomposition matrices into each layer of the Transformer architecture, reducing the number of trainable parameters. Quantization improves over LoRA by quantizing the transformer model to 4-bit precision.The number of trainable parameters for QLoRA is 3.4 million.

**Training command:** For QLora training, we apply the command
CUDA_VISIBLE_DEVICES=0 llamafactory-cli train examples/lora_single_gpu/llama3_lora_sft.yaml.
For Full training, we apply the command
CUDA_VISIBLE_DEVICES=0,1,2,3 python -m torch.distributed.run \
--nproc_per_node $NPROC_PER_NODE --nnodes 1 --standalone \
src/train.py full_fine_tunning/single_node.yaml
In the commands, CUDA_VISIBLE_DEVICES specifies how many GPUs to use within a compute node, and the python and yaml files can be found in LLama-Factory Github.

Detailed parameters and configurations used are:

| ***Hyper-parameters*** | ***Full Fine-Tuning*** | ***QLoRA Fine-Tuning*** |
|---|---|---|
| Learning Rate | 5e-6 | 1e-4 |
| Fine-tuning Type | Full | Lora |
| Quantization Bit | NA | 4 bits |
| Per_device_train_batch_size | 16 | 16 |
| Gradient_accumulation_steps | 8 | 8 |
| Training_epochs | 6 | 15 |
| Lr_scheduler_type | cosine | cosine |
| Warmup_steps | 0.05 | 0.05 |
| Distributed Training | Deepspeed | NA (Trained on a single GPU) |
| Optimizer | adamw_torch | adamw_torch |
| Dataset | FinalQA | FinalQA |

**Table**. Training details for our instruction tuning experiments. Both Full Fine-Tuning model and QLoRA-Tuning model are trained based upon LLama3-8B-Instruct model with Data FinalQA. The QLoRA Fine-tuning model costs a single A100 GPU with 1hr, while the Full Fine-tuning model is trained on 4 A100 GPUs for 5hrs. Each GPU has memory of 80G.

**Choice of epoch number:** During training, we varied the number of training epochs and found that finetuning >15 epochs does not help. In particular, full-parameter fine-tuning for 20 epochs did not improve the performance in gene-editing questions compared to CRISPR-Llama3 trained with 6 epochs. We tested it for multiple-choice questions[65] in the FinalQA dataset, and it attained a score of 90% that was only comparable to CRISPR-Llama3 (91%). What's more important is that fine-tuning for >15 epochs actually degrades the model's performance on general questions, due to over-optimization/overfitting to the small dataset used for finetuning.

Our choice of epoch number and observation of overfitting is consistent with common practice in LLM research. In general LLM research, while the base model is often pretrained on large amount of data entries using a large number of epochs, finetuning usually takes only a few epochs (2-15 epochs). The reason is that finetuning a model on a small domain specialized dataset could easily cause overfitting and catastrophic forgetting. A model that is "over-finetuned" could appear to memorize the dataset used for finetuning but it cannot generalize the knowledge and even forget common senses learnt via pre-training. The LIMA paper[70] suggests that supervised fine-tuning (SFT) only requires a small demonstration dataset. In their setting, they finetuned llama with 15 epochs with 1000 curated (question, response) pairs and showed remarkable performances. The BERT paper[71] finetuned its model using only 2-3 epochs, and the RoBERTa paper[72]finetuned its model using 10 epochs.

### 6. Evaluation and Rubrics

To evaluate the fine-tuned models, we compiled an independent testset comprising three sets of questions:

1. **Multi-choice Questions**: 20 multiple-choice questions curated from two sources of online knowledge exams (https://worldscienceu.com/quizzes/2-3-test-crispr-knowledge and https://quizizz.com/admin/quiz/5e977345a5b8a8001fe3478e/crispr-quiz), designed to test the model's ability to distinguish correct from incorrect answers related to gene-editing, including both basic fact-checking and experimental design questions.
2. **Basic Knowledge QA**: A set of 10 questions assessing the model's understanding of fundamental CRISPR knowledge, from the UC Berkeley Innovative Genomics Institute's online CRISPR FAQ[60].
3. **Real-world Problem Solving (STAR_QA)**: A curated set of 10 open-ended questions, published by the journal STAR Protocols[61], reflecting real-world challenges encountered by scientists during CRISPR gene-editing experiments.

For each multiple choice question, we generated 10 answers using the models and then the average scores of the questions were collected for scoring. The scoring followed a stringent metric: the model will get a score of 1.0 for the question only when the model was able to correctly answer all the keys, otherwise it will get a score of 0.0. Finally all scores were calculated to yield the average score of each model. For each question in "Basic Knowledge QA" and "Real-world Problem Solving", we generated 2 independent answers using the 3 models for all the open-ended questions, including the basic knowledge and the real-world problem-solving questions. We then asked three independent human experts in gene-editing to evaluate the answers to these questions. The scoring rubrics for open-ended questions by human evaluations are: Score of 1 if the response is mostly correct and useful. Score of 0.5 if the response has errors but still helpful. Score of 0 if the response is not correct at all and not useful.

**Fine-tuning evaluation findings:** Evaluation results of the fine-tuned LLM can be found in **Supp. Figure 1.** The fine-tuned model outperforms the baseline un-finetuned model on simple multiple choice questions by a moderate 8% and on real-world research questions by ~20% (**Supp. Figure 1c).** The fine-tuned model's improved performance on open-ended problem-solving questions showed that our instruction tuning improved the capability of the model for answering domain-specific questions for gene-editing. The fine-tuned model provided helpful, expert-like suggestions for questions like "What is a good negative control guide RNA for gene-editing?" and "When I perform CRISPR experiment, my cells keep dying after single-cell sorting. Any advice on how to troubleshoot these issues?" (**Supp. Figure 3**). These results highlight the benefit of using domain expert discussions to improve LLM performance on scientific problems.

### 7. Detailed Methodology of LLM Fine-tuning

**Full Instruction Fine-tuning:** Instruction fine-tuning involves training a large language model (LLM) to perform well on tasks where it follows user instructions. It's a process where the model is fine-tuned on labeled datasets, where each input corresponds to a specific desired output. The goal is to align the model's behavior with the human expert's intent. An LLM (e.g. Llama) is typically a neural network based on the transformer architecture[66]. Let the model be parameterized by $\theta$, and given an input question $x$, the model outputs a probability distribution $P_\theta(y|x)$ over the possible outputs $y$. During instruction fine-tuning, the model is trained on pairs $(x_i, y_i)$, where $x_i$ is the *question (+ context)* in FinalQA and $y_i$ is the *answer*. The goal of instruction fine-tuning is to minimize the difference between the model's predicted output and the true output (ground truth) for a given instruction. The standard loss function used is the **cross-entropy loss**, which measures how well the predicted probability distribution $P_\theta(y_i\,|x_i\,)$ aligns with the actual distribution. The cross-entropy loss for a single instruction-output pair $(x_i, y_i)$ is defined as:

$$L(\theta; x_i, y_i) = -\sum_{t=1}^{T} \quad log\ P_\theta(y_{i,t}|x_i, y_{i,<t})\ ,$$

where $y_{i,t}$ is the token at position $t$ in the output sequence $y_i$, $T$ is the length of the output sequence,$P_\theta(y_{i,t}|x_{i,y_{i,<t}})$ is the probability assigned by the model to the token $y_{i,t}$, conditioned on the input $x_i$ and all previously generated tokens $y_{i,<t}$. For the FinalQA dataset of instruction-output pairs $\{(x_i, y_i)\}^N{}_{i=1}$, the total loss is: $L(\theta) = \frac{1}{N}\sum_{i=1}^{N} \quad L(\theta; x_i, y_i)$. This loss function encourages the model to assign higher probabilities to correct outputs (i.e., $y_i$) for a given input instruction $x_i$.

To minimize the loss, we update the model parameters using gradient descent. The parameter update rule at step $t$ is given by: $\theta_{t+1} = \theta_t - \eta\nabla_\theta L(\theta_t)$, where the learning rate $\eta$ is a hyperparameter. In practice, it is common to use **AdamW optimizer**[67] (a variant of gradient descent) in fine-tuning tasks. It adjusts the learning rate based on past gradients, making it more efficient for training large models.

**QLoRA Fine-tuning:** We first explain LoRA[68] technique then introducing quantization. LoRA (Low-Rank Adaptation) is a technique for fine-tuning large language models (LLMs) that reduces the number of trainable parameters, making fine-tuning more efficient. Instead of updating all the parameters of the LLM, LoRA introduces low-rank matrices to adapt pre-trained models, considerably reducing the computational and memory overhead. LoRA assumes that weight updates during fine-tuning lie in a low-rank subspace. Instead of directly updating the large weight matrices of the model, LoRA approximates these updates with low-rank matrices. The original large weight matrices are kept frozen, and only the small low-rank matrices are updated.

Let $W_0 \in R^{d\times k}$ represent a pre-trained weight matrix of the LLM, where $d$ is the input dimension and $k$ is the output dimension. In *standard fine-tuning*, we would update $W_0$ directly, i.e., $W = W_0 + \varDelta W$, where $\varDelta W$ is the full-rank weight update matrix learned during fine-tuning. In *LoRA fine-tuning*, instead of learning the full-rank matrix $\varDelta W$, one could decomposes it into two low-rank matrices: $\varDelta W = AB^\top$, where: $A \in R^{d\times r}, B \in R^{r\times k}$, where $r$ is much smaller compared to $d$ or $k$. This means that during fine-tuning, we are learning the matrices $A$ and $B$, both of which have much fewer parameters compared to $W_0$. The updated weight matrix becomes: $W = W_0 + AB^\top$.

*Loss Function.* For LoRA fine-tuning, it also uses the cross-entropy loss. For an input $x$ and its corresponding label $y$, the trainable parameters are low-rank matrices $A, B$ with the loss $L(A, B; x, y) = -\sum_{t=1}^{T} \quad log\ P_\theta(y_t|x, y_{<t})$ . The difference is that instead of optimizing the full $W$, we are now optimizing the low-rank matrices $A$ and $B$.

*Gradient update.* The goal is to minimize the loss with respect to $A$ and $B$. The gradient update for LoRA also follows the gradient descent mechanism (with $\eta$ being the learning rate):

$$A_{t+1} = A_t - \eta\nabla_A L(A_t, B_t; x, y);\ B_{t+1} = B_t - \eta\nabla_B L(A_t, B_t; x, y).$$

Since $A$ and $B$ are much smaller than $W_0$, the computational cost is considerably reduced.

*Quantization.* QLoRA[69] extends LoRA by applying quantization to the frozen pre-trained weights in order to further reduce memory usage. The model weights are quantized into lower-precision formats (e.g., 4-bit), which allows for loading much larger models into memory. At the same time, QLoRA retains LoRA's low-rank adaptation for efficient fine-tuning.

#### C. Gene-Editing-Bench testset and full evaluation procedures

### 1. Gene-editing experiment planning evaluation

To assess the capability of the LLM-planner in autonomously generating a list of subtasks based on user requests, we developed a gene-editing experiment planning test set comprising 50 typical user queries spanning various gene-editing scenarios. Experts in the CRISPR field curated and labeled the ground truth subtask lists for each user request.

We generated three independent batches of answers using CRISPR-GPT and baseline models. The generated subtask lists were compared with the expert-labeled ground truth subtask lists. For each request, we defined tasks present in both the generated and ground truth lists as **true positives**, tasks appearing only in the generated list as **false positives**, tasks present in the ground truth list but missing in the generated list as **false negatives**, and tasks absent from both lists as **true negatives**.

To quantify performance, we calculated overall accuracy, precision, recall, and F1 scores using the following standard formulas:

- **Accuracy** = *Accuracy = (TP + TN) / (TP + TN + FP + FN)*
- **Precision** = *Precision = TP / (TP + FP)*
- **Recall** = *Recall = TP / (TP + FN)*
- **F1 Score** = *F1 Score = 2 × Precision × Recall / (Precision + Recall)*

Additionally, to evaluate whether the generated subtasks were presented in the correct order according to the user request, we utilized the **Levenshtein distance (Ld)** between the generated and ground truth sequences of subtasks. Each subtask was treated as a unique symbol, with costs incurred for additions, deletions, or substitutions (cost = 1). We report the **average normalized Levenshtein distance (Ldn)** as:

- *Ldn = Ld / (n × N)*

where **n** is the total number of subtasks per user request and **N** is the total number of user requests.

Simultaneously, we generated three independent batches of answers using **gpt-4o** and **gpt-3.5-turbo** on the same set of user requests. Gene-editing experts reviewed and labeled each response as either correct or incorrect. The accuracy for each model was calculated as:

- *Accuracy = Number of Correct Responses / Total Number of User Requests*

### 2. Gene-editing delivery selection evaluation

The choice of delivery is a critical step of designing a successful gene-editing experiment. There's a saying in the field that "the challenges of gene-editing is delivery, delivery, delivery". Even in a laboratory research setting, there are often many possible choices for delivering the CRISPR system into target cells of interest. If one searches online via Google or general LLMs, the most likely results would be a list of possible options, such as chemical transfection, liposome-based transfection, electroporation/ nucleofection, viral-based delivery, lipid nanoparticle delivery, etc. We propose that LLM agents are uniquely positioned to address the complexities given its large knowledge base and ability to perform logical reasoning in a defined domain like CRISPR gene-editing. In CRISPR-GPT, we designed the LLM agent with a series of expert instructions, and ability to use external tools such as performing web and literature search and ranking, allowing it to complete the task like a human expert.

To evaluate the effectiveness of our delivery suggestion function, we created a delivery selection dataset, which includes 50 typical user queries regarding the selection of CRISPR delivery methods across various biological systems. For each query, we engaged multiple CRISPR experts to assess the applicability of six common CRISPR delivery methods:

- a. Plasmid Transfection
- b. Lentivirus/Retrovirus
- c. RNP/mRNA Electroporation
- d. RNP/mRNA Microinjection
- e. mRNA LNP
- f. AAV

Each method was scored on a scale from 0 to 2, with 2 indicating the most suitable and commonly used method, 1 indicating potential usability under special conditions, and 0 suggesting infeasibility or rare use. Following the evaluation, experts convened to consolidate their assessments into a unified score sheet, which served as the ground truth for the test sets.

Subsequently, three independent batches of responses were generated using CRISPR-GPT (with/without the "literature search" function), gpt-3.5-turbo, and gpt-4-turbo. Each model was prompted to propose a primary and a secondary delivery method for each query. Responses were then evaluated against the ground truth, with the primary delivery method assigned a weight of 2 and the secondary a weight of 1. Scores for all requests were summed and percentage correctness reported for each category of request.

#### 3. guideRNA design evaluation

To evaluate the performance of the sgRNA design function, we constructed the gRNA design benchmark dataset, consisting of 50 typical user queries related to sgRNA design for CRISPR-mediated knockout, activation, or interference. To address these queries, we implemented four key functions — SELECT, BETWEEN, ORDERBY, and TOP — to process predesigned sgRNA tables and retrieve relevant sgRNA information for presentation to users. Experts in the CRISPR field manually curated a list of functions and their corresponding parameters for each user query, validating them to serve as the ground truth.

We then prompted CRISPR-GPT to generate three independent batches of function lists and relevant parameters from user queries. The generated responses were compared to the ground-truth answers, and we calculated the accuracy of both the functions (i.e., correct function selection and correct order) and the parameters (i.e., correct parameters per function).

Simultaneously, we prompted gpt-4o and gpt-3.5-turbo to generate three independent batches of function lists and corresponding parameters. CRISPR experts reviewed these responses and labeled each function list and its corresponding parameters as either correct or incorrect, based on whether the proposed functions were relevant, in the correct order, and whether all relevant parameters were accurately captured. The accuracy of both the function (per request) and the parameters (per function) was then calculated.

#### 4. Gene-editing QA evaluation

To assess the performance of the QA mode of CRISPR-GPT, we developed the Gene-editing QA benchmark dataset, which includes 138 questions covering a broad range of gene-editing topics. These topics encompass CRISPR basic knowledge, experimental troubleshooting, CRISPR applications, ethics, and safety. The full testset expands on the previous smaller set used for fine-tuning and was sourced from real-world CRISPR inquiries compiled from public sources and human experts. All questions were then carefully filtered by human gene-editing experts to eliminate errors and inconsistencies.

For the evaluation of QA mode, we selected 31 representative questions from the testset and prompted CRISPR-GPT, gpt-3.5-turbo, and gpt-4o to generate responses. The responses for each question were anonymized, and three CRISPR experts were asked to evaluate and score the answers across four key aspects: accuracy, reasoning, completeness, and conciseness (detailed rubrics are in **C6** below). The scores from this fully blinded evaluation by experts were averaged to calculate the final performance scores.

In the evaluation, human evaluators observe that general-purpose LLMs sometimes make factual errors and tend to provide long answers that are not all relevant to the questions (**Figure 4f**). For example, for the question "*Why doesn't Cas9 cleave the original CRISPR sequence in the bacterial genome?*", GPT4o gave the correct answer (PAM) but also a factually wrong, non-relevant reason (crRNA mismatch), while our QA Mode gave a precise answer (**Ext. Data Figure 2a**). In another example, for the question "*What's the difference between Cas9 and Cas12a?*" CRISPR-GPT gave a concise and correct answer. However, GPT4o gave a long list of differences but incorrectly claimed that Cas9's multiplexing ability is an advantageous feature over Cas12a. In fact, Cas12 is the better system for multi-target gene editing (**Ext. Data Figure 2b**). For a third example, let's look at a question about solving cell growth issues in an experiment where a scientist performed Cas9 editing followed by single-cell sorting using MCF-7 cells. For this question, CRISPR-GPT QA Mode provided a fully accurate summary of potential reasons and actionable solutions. In contrast, GPT-4o's responded with a long list of 9 itemized factors/options, but at least 2 of them are not applicable to MCF-7 cells (**Ext. Data Figure 2c**). Overall, evaluation results confirmed that the multi-source QA Mode in CRISPR-GPT is better at answering advanced research questions about gene-editing.

#### 5. Human user experience evaluation

To evaluate user observations of CRISPR-GPT across various tasks, we invited 8 independent CRISPR experts to test the web-based CRISPR-GPT agent. Each expert was asked to test two gene-editing requests using Meta mode and two gene-editing requests using Auto mode.

For Meta mode, experts were tasked with designing two gene-editing requests and scoring their experience for each task across four aspects: accuracy, reasoning and action, completeness, and conciseness, using a scale of 1 (Poor) to 5 (Excellent) (details in Supp. Note **C7** below). Experts also tested gpt-3.5-turbo and gpt-4o using equivalent prompts and the same scoring criteria via OpenAI APIs (so blinded to the version of models). At the end, experts provided an overall score and comments.

For Auto mode, experts tested two different gene-editing requests with CRISPR-GPT, gpt-3.5-turbo, and gpt-4o, scoring each model using the same rubric. All scores were summarized and averaged.

### 6. QA Mode evaluation rubrics

**Accuracy**

- 1 (Poor): The answer contains multiple factual errors or shows a misunderstanding of CRISPR technology.
- 2 (Fair): The answer has some correct elements but also includes substantial inaccuracies.
- 3 (Average): The answer is mostly accurate but may contain minor errors or oversights.
- 4 (Good): The answer is accurate, with only negligible errors that do not impact the overall validity of the information provided.
- 5 (Excellent): The answer is completely accurate, reflecting the current state of CRISPR knowledge.

**Reasoning**

- 1 (Poor): The reasoning behind the answer is flawed or nonexistent; the logic is unclear or incorrect.
- 2 (Fair): The answer provides a rationale, but it is weak and may not support the conclusion or design effectively.
- 3 (Average): The answer's reasoning is solid for the most part, with some areas that could be better supported or explained.
- 4 (Good): The answer provides strong reasoning with clear and logical support for all claims and suggestions made.
- 5 (Excellent): The answer's reasoning is exceptional, providing insightful, well-supported explanations that enhance understanding of CRISPR knowledge.

**Completeness**

- 1 (Poor): The answer is incomplete and lacks critical information required to form a complete understanding.
- 2 (Fair): The answer covers some necessary points but omits several important aspects that would be needed.
- 3 (Average): The answer is fairly comprehensive but could be improved with additional details or coverage of more nuanced aspects.
- 4 (Good): The answer is thorough, covering nearly all aspects required for a complete understanding and successful experimental setup.
- 5 (Excellent): The answer is entirely comprehensive, leaving no question unanswered and providing a full suite of information needed.

**Conciseness**

- 1 (Poor): The answer is overly verbose and contains much irrelevant information, making it difficult to extract useful insights.
- 2 (Fair): The answer is longer than necessary with some extraneous content but still delivers a fair amount of relevant information.
- 3 (Average): The answer conveys the necessary information with some unnecessary detail but remains clear and understandable.
- 4 (Good): The answer is concise, with well-organized content that is directly relevant to the question asked, without any unnecessary information.
- 5 (Excellent): The answer is exceptionally concise, communicating the required information efficiently and effectively.

### 7. User experience evaluation rubrics

**Accuracy**

- 1 (Poor): The answer contains multiple factual errors or shows a misunderstanding of CRISPR technology.
- 2 (Fair): The answer has some correct elements but also includes substantial inaccuracies that could lead to flawed experimental design if followed.
- 3 (Average): The answer is mostly accurate but may contain minor errors or oversights.
- 4 (Good): The answer is accurate, with only negligible errors that do not impact the overall validity of the information provided.

- 5 (Excellent): The answer is completely accurate, reflecting the current state of CRISPR research and methodologies.

**Reasoning and Action**

- 1 (Poor): The reasoning behind the answer is flawed or nonexistent, and the model fails to perform relevant actions. There is no logical connection between the reasoning and any actions attempted.
- 2 (Fair): The reasoning is present but weak, with limited support for the conclusions or actions taken. The model attempts to perform actions, but they are either incomplete or not well-aligned with the problem at hand.
- 3 (Average): The reasoning is mostly solid, though there are areas that could be better explained or supported. The model performs appropriate actions but lacks precision or optimal efficiency in its execution.
- 4 (Good): The reasoning is clear and well-supported, providing logical justification for the actions taken. The model performs the actions effectively, demonstrating a good alignment between reasoning and execution.
- 5 (Excellent): The reasoning is exceptional, offering deep insights and clear explanations. The model performs actions flawlessly, demonstrating innovation, precision, and effectiveness in executing the tasks based on the reasoning provided.
-

**Completeness**

- 1 (Poor): The answer is incomplete and lacks critical information required to form a complete understanding.
- 2 (Fair): The answer covers some necessary points but omits several important aspects that would be needed for a thorough CRISPR design.
- 3 (Average): The answer is fairly comprehensive but could be improved with additional details or coverage of more nuanced aspects of the design.
- 4 (Good): The answer is thorough, covering nearly all aspects required for a complete understanding and successful experimental setup.
- 5 (Excellent): The answer is entirely comprehensive, leaving no question unanswered and providing a full suite of information needed for CRISPR experimental design.

**Conciseness**

- 1 (Poor): The answer is overly verbose and contains much irrelevant information, making it difficult to extract useful insights.
- 2 (Fair): The answer is longer than necessary with some extraneous content but still delivers a fair amount of relevant information.
- 3 (Average): The answer conveys the necessary information with some unnecessary detail but remains clear and understandable.
- 4 (Good): The answer is concise, with well-organized content that is directly relevant to the question asked, without any unnecessary information.
- 5 (Excellent): The answer is exceptionally concise, communicating the required information efficiently and effectively.

### D. Limitations, safety and ethical consideration, dual-use study

**1. Limitations of current study**

Here we discuss several limitations with the LLM agent described in this work for gene-editing and related biological experiments. First, while CRISPR-GPT can effectively design individual components such as guide RNAs and primers, additional connection with latest advances in genome/protein foundation models, plasmid design tools, and other machine learning models, could enable design tasks beyond gene-editing, e.g. design of tailored therapeutic molecules like mRNAs. Second, the agent's performance may be limited in complex gene editing requests or rare biological cases that are not well-represented in its training data or knowledge base. Continual updating of CRISPR-GPT's domain knowledge, safeguards, and expanding its integrated tool sets will be important to address increasingly sophisticated applications. Third, the real-world validation of CRISPR-GPT, though promising, may not fully encompass the diversity and complexity of gene-editing applications across different organisms or cell types. Further testing and refinement will improve its reliability across a wider range of experimental conditions. Overall, regular auditing and updating of the agent's modules in line with the latest scientific and regulatory developments will help to bring exciting applications and responsible uses of genome engineering technologies.

**2. Implications for agents towards broader categories of biological experiments**

The LLM agent in the current study is designed for a major, but specialized type of biological experiment, CRISPR gene-editing. While the current agent serves a niche area, the challenges and difficulties we observed, as well as the solution and approach we proposed, have the potential to transfer to other areas of biological experiments and research topics.

First, the human-AI collaborative approach demonstrated in CRISPR-GPT, where the LLM agent works alongside researchers to design experiments, could also have far-reaching implications. The LLM was designed through mimicking aspects of the thought processes of human domain experts, while also leveraging "Chain-of-thought" prompting and "state machine / memory" architecture that are state-of-the-art advance in LLM engineering, thus CRISPR-GPT showcase that optimal solution may require the best of both science and AI worlds. By leveraging the strengths of both human expertise and artificial intelligence, this paradigm has the potential to accelerate discovery and innovation across various biological disciplines. Further integration with additional LLM agents such as those assisting researchers in hypothesis generation, data visualization, and even the interpretation of results, could ultimately lead to more efficient and effective research processes.

Second, the modular architecture and task decomposition strategy employed in CRISPR-GPT could serve as a blueprint for developing LLM agents in other areas of biological research. Breaking down the experimental design process into discrete, manageable tasks and implementing them as interconnected state machines allows for a structured, systematic approach to problem-solving. This modular framework also facilitates the incorporation of new tools, datasets, and experimental techniques as they emerge, ensuring the agent remains up-to-date with the latest advancements in the field.

Third, one key aspect of CRISPR-GPT that could be broadly applicable is the integration of domain-specific, curated knowledge and external tools into the LLM-based agent. By equipping the agent with curated biological databases, protocols, and computational tools tailored to a particular field of biology, researchers can leverage the reasoning capabilities of LLMs to navigate complex experimental design tasks across various domains. We expect this will apply to additional areas, such as protein engineering and directed evolution, metabolic pathway optimization, or high-throughput screening assays.

However, the development of LLM agents for broader categories of biological experiments will also require addressing the limitations and challenges highlighted in the CRISPR-GPT study. These include the need for robust fact-checking and validation mechanisms to mitigate the risk of hallucinations, regular updates to the agent's knowledge base and ethical/safety modules, and the development of more advanced natural language processing capabilities to handle the complexity and diversity of biological terminology and concepts. As the field of AI continues to advance, the lessons learned from CRISPR-GPT will undoubtedly inform the design and implementation of LLM agents across a wide range of biological research areas. By embracing these insights and adapting them to the unique challenges of each domain, we can harness the power of language models to revolutionize the way we conduct scientific research, ultimately leading to groundbreaking discoveries and transformative applications in biology and beyond.

**3. Dual-use study for safety and ethical considerations**

Substantial concerns exist regarding the use of gene-editing methods: (**1**) **Heritable human edits**, there are ethical concerns and societal risks to edit human genome that could lead to heritable alterations to the common genetic pool of the human species[1]; (**2**) **Pathogen engineering**, the use of gene-editing to engineer pathogenic organisms such as highly dangerous viruses is an important biosafety risk[2].

Our implementation of CRISPR-GPT agent have 2 layers of protection / prevention:
**Layer 1**: keywords filtering during prompt / request. Specifically, for layer 1, a list of keywords are screened, as listed in **Supplementary Material 1.**
**Layer 2**: addition of explicit warning and consenting step

To assess the risks associated with inappropriate usage of CRISPR-GPT agent, we designed a set of experiment requests for biological targets of concern, covering the above areas. We submitted these requests as prompts to the agent. We then examine the output of CRISPR-GPT to determine how the agent may or may not proceed with these requests (**Ext. Data Figure 3**).

**4. Protection of user genome data privacy:**
A notable unique feature of biological experiment is the potential of involving human genome data with considerable privacy and societal implications. There is growing concern that online biological tools would exploit sequence information that could be identifiable. For this part, a set of guidelines should be followed, such as the Health Insurance Portability and Accountability Act (HIPAA) in the US. Thus, we set up a hard-coded recognition of identifiable nucleic acid sequences at the core of CRISPR-GPT. Once recognized, under no circumstances would sequence information be passed to the LLM agent, blocking any possible leakage of user sequence data (**Ext. Data Figure 3**).

**5. Concluding note on ethical, safe usage of LLM agent for biological experiments**
Taking together our observations from tests, the results from the dual-use study, and the privacy considerations, we believe that our work demonstrates the importance of having a set of safety guardrails and privacy protection mechanisms for biological LLM agents. This is to ensure responsible and secure usage of these agents for designing biological experiments. Specifically, we have the following recommendations:

1. Explicit rejection of requests through stringent, explicit logic, with back-end prompt engineering to avoid any risk from explicit or concealed requests
2. Zero tolerance for the storage or transmission of user supplied sequence data to the underlying LLM, hard-coded to ensure that this is required regardless of the API or other types of interface being used.
3. Implementation of real-time updates to continuously monitor new technology development and data, ethical risks, as well as guidelines from WHO, IGSC, and the community.
4. Security measures such as the authentication mechanism we are using to avoid any potential attack, bypass, or modification to the underlying LLM agent.
5. Policy awareness should also be part of the consideration when developing any LLM agent for biological experiments, in line with international and national governing body guidelines and regulations, such as the Global guidance framework for the responsible use of the life sciences: mitigating biorisks and governing dual-use research (WHO).

The above dual-use study and examples were performed in a purely computational manner by researchers. None of the examples listed were implemented in actual experiments. We reiterate that, under no circumstances should any individual or organization attempt to perform gene-editing that could lead to heritable changes or germline cell alterations in humans or perform genetic engineering of any dangerous pathogens. Specific list of pathogens are exemplified by the International Gene Synthesis Consortium (IGSC) in the IGSC's Harmonized Screening Protocol "Regulated Pathogen Database", which is assembled and curated by the IGSC to include data from all organisms on the US Select Agent and Toxin list (https://www.selectagents.gov/sat/list.htm), the Australia Group Common Control List (https://www.dfat.gov.au/publications/minisite/theaustraliagroupnet/site/en/controllists.html), and other national lists of regulated pathogens and toxins.

### E. Agent Prompt Formats

**1. The prompt format of the LLM Planner Agent.**

We use the following prompt format for task decomposition for the LLM planner in the automation mode. The LLM planner interprets the user's request and decomposes it into a list of tasks. The LLM planner is prompted to respect the task dependencies stated in the Task Description Table.

Please act as an expert in CRISPR technology. Given the user input, think step by step and generate a list of tasks for execution. First refer to the task description table below, and try to figure out if the user needs to directly jump into a task, or the user needs to complete several tasks. Make sure to respect the task dependencies and include all dependent tasks in the list.

Please format your response and make sure it is parsable by JSON.

#### Task Description Table
{Task Description Table}

#### Demonstrations:
If the user only needs to design guideRNA for knockout, then return ['knockout. StateStep1', 'knockout.StateStep3']. Reason: this directly matches knockout. StateStep3. But it needs to complete knockout.StateStep1 first, so both 'knockout. StateStep1' and 'knockout.StateStep3' are returned.

User Input:
"{user_message}"

Response format:
{{
"Thoughts": "",
"Tasks": ["", ""] ## a list of task names
}}

The task description table contains all the implemented tasks and their dependencies. See example below:

For knockout task name: task descriptions: dependency
knockout.StateStep1: Cas System selection for knockout : none
knockout.StateStep2: Delivery approach selection for knockout : none
knockout.StateStep3: guideRNA design for knockout : needs to complete knockout. StateStep1 first

**2. The prompt format of the User-proxy Agent.**

We synthesize the relevant information into {system_message}, including the instruction of the current state, the interaction history between the agent and the system, and potentially the results from external tools and libraries. Next, we supply the meta request of the user in {meta_prompt}. Then we prompt the LLM-agent to understand the current state and make decisions on behalf of the user.

Please act as you are using the CRISPR design tool. Given the user meta request, the current inquiry provided by the tool, think step by step and generate an answer to the questions. Please format your response and make sure it is parsable by JSON.

Rules:
1. Answer the inquiry directly on behalf of the user. Don't raise any additional questions to the user.
2. If the inquiry is a multiple-choice question, then directly output one choice.
3. If the inquiry asks you to supply any gene sequence, then answer the question with "I don't know" and let the user take manual control.

User Meta Request:
"{meta_prompt}"
Current Inquiry:
"{system_message}"
Response format:
{{

"Thoughts": "",
"Answer": ""
}}

### F. Example publications utilized in RAG system
